# Visualization of Bluetongue virus RNA segment networks in infected cells: multipartite genomic RNA assortment is independent of viral proteins NS2 and VP6

**DOI:** 10.64898/2025.12.04.692307

**Authors:** Dong-Sheng Luo, Po-Yu Sung, Polly Roy

## Abstract

Bluetongue virus (BTV), with a genome of ten double-stranded RNA segments (S1–S10), is an emerging animal pathogen causing major economic losses in livestock worldwide. BTV replication involves RNA-RNA and RNA-protein interactions, with RNA-binding proteins, VP6 and NS2 playing key roles in genome assembly and RNA packaging. To explore the dynamics of RNA segment interactions and the roles of VP6 and NS2 in RNA complex formation, we used RNA fluorescence *in-situ* Hybridisation Chain Reaction (HCR), along with site-specific mutagenesis and reverse genetics. We found that RNA segments interact sequentially, from the smallest (S10) to the largest (S1), forming a single complex that includes the entire genome. This process is independent of VP6 or NS2, although NS2 enhances the assembly of larger segments. Additionally, we show that VP6 binds to +ssRNAs before their incorporation into viral assembly factory (inclusion bodies/VIBs). These findings reveal that RNA-RNA interactions, rather than primary replicase proteins, govern the sorting and recruitment of genome segments. Our data offer new insights into BTV RNA packaging, showing that genome segments destined for packaging and dsRNA synthesis are segregated through complex formation, distinct from +ssRNAs used in protein synthesis, including those encoding the replicase complex.

## INTRODUCTION

Bluetongue virus (BTV), a significant and emerging animal pathogen, is the prototype of the *Orbivirus* genus (22 species; and more than 130 serogroups) within the subfamily *Sedoreoviridae of Reovirales* (16 distinct genera) (1). *Reovirales* members include viruses that infect a wide range of vertebrates, including humans and animals, and plants. A unique feature of the *Reovirales* members is that their genomes are made up of double-stranded RNA segments (dsRNAs, 9-12 segments). All members are nonenveloped viruses with icosahedral capsids, made up of concentric protein layers. To understand how dsRNA segments are differentiated for selection and packaging from host RNAs within the *Reovirales*, we used BTV as model virus. The BTV genome is comprised of 10 dsRNA segments, ranging in size from 3.95 kb to 0.8 kb, grouped into three size classes: large, medium, and small (2, 3). It encodes seven structural proteins (VP1-VP7) and four non-structural proteins (NS1, NS2, NS3/NS3A and NS4), with the structural proteins forming a complex icosahedral virion consisting of an outer capsid (VP2, VP5) and an inner core (VP1, VP3, VP4, VP6, and VP7) (3–7). The final product of BTV disassembly is the transcriptionally active core particle in which the resident RNA polymerase has been activated. The ten genomic dsRNA segments are transcribed simultaneously and repeatedly to generate ssRNAs, each of which is released into the cytoplasm through discrete channels at the core apices (8). These transcripts serve as mRNAs for viral protein synthesis but later also as ssRNA templates for progeny dsRNA synthesis. Of the five core proteins, the three minor proteins, VP1 (polymerase), VP4 (capping enzyme), VP6 (RNA packaging protein) and the major protein VP3 (forming the innermost layer) together with the NS2 protein form the primary replicase complex (9, 10). NS2 is a phosphorylated protein and phosphorylation of NS2 is responsible for the formation of viral inclusion bodies (VIBs) that serve as the virus assembly sites in the cytoplasm. NS2 has a strong affinity for single-stranded RNA (ssRNA) (11).

During BTV replication, newly synthesized transcripts (+ssRNA) segments are first packaged into assembling viral cores, which subsequently act as templates for synthesizing genomic dsRNA segments (12). Previous *in vitro* evidence indicated that BTV RNA packaging is driven by the formation of complex RNA networks, a process which follows a specific sequence, starting with the smallest segment, S10, which initiates RNA-RNA interactions with other small segments to form an initial complex (S7-S10). Medium-sized (S4-S6) and larger ssRNAs (S1-S3) are then progressively recruited, resulting in the complete genome complex being packaged into the capsid (13–15). Although NS2 is not necessary for *in vitro* ssRNA complex formation or core assembly (12, 13, 15), it is essential for virus replication in infected cells (9, 16). Reverse genetics (RG) studies demonstrated that both NS2 and VP6 are essential for BTV replication as modified BTV strains lacking NS2 or VP6 failed to replicate in normal cells but were able to propagate in NS2- or VP6-expressing helper cell lines (16, 17). Moreover, when VP6-deficient viruses are grown in complimentary helper cells and used to infect normal cells, viral proteins are synthesized and assembled into empty particles lacking the viral genome (16). Despite these findings, the precise functions of NS2 and VP6 in virus replication remain unclear. While both are believed to play critical roles in RNA recruitment and packaging in infected cells, it is also hypothesized that they may have additional functions, such as facilitating the packaging of viral RNA into the capsid during assembly.

To examine the mechanism of BTV RNA recruitment and packaging, we adopted an RNA fluorescence *in-situ* Hybridisation Chain Reaction (HCR) technique tailored for BTV RNAs (13). HCR (18–20), is an isothermal nucleic acid hybridisation signal amplification technique that has gained significant attention due to its enzyme-free operation and suitability for ambient conditions. It is an alternative to traditional RNA Fluorescence *in-situ* Hybridisation (FISH) and has been increasingly adopted for a broad range of applications. Its enhanced sensitivity, enzyme-free amplification, and adaptability make it particularly effective for viral RNA detection and localization (21–24) and it been used successfully to image viral RNAs (25–28). We demonstrated elsewhere its high sensitivity and specificity in localizing transfected BTV viral RNAs within BSR cells (13). HCR ensures that multiple probes bind simultaneously to the targeted BTV RNA segments, thereby producing high-intensity fluorescence signals and enabling precise localization of viral RNAs within infected cells. Using this technique together with site-specific mutagenesis and reverse genetic-based virus recovery, we demonstrate for the first time in BTV-infected cells how BTV RNA +ssRNAs segments are selected for packaging and what roles are played by VP6, the RNA packaging protein and NS2/VIBs, the virus assembly protein and assembly site. This study of RNA-RNA interaction in infected cells, RNA-protein interaction and larger RNA complex formation provides valuable insights into the BTV assembly pathway and RNA recruitment/packaging mechanisms. Moreover, it offers a foundation for exploring assembly processes in other dsRNA viruses, helping to determine the sequential order of genome packaging and core/capsid formation.

## MATERIALS AND METHODS

### Cell, virus, plasmid, mutagenesis and antibody

BSR (BHK-21 cells subclone, ATCC^â^ CCL10^TM^), BSR-NS2 (BSR cells stably expressing NS2/segment 8 of BTV) and BSR-VP6 (BSR cells stably expressing VP6/segment 9 of BTV) cells were cultured in Dulbecco’s Modified Eagle Medium (DMEM) (Sigma) supplemented with 10% fetal bovine serum (FBS, Sigma) at 37 °C with 5% CO_2_. Medium for BSR-NS2 and BSR-VP6 were also supplemented with puromycin (7.5 µg/ml). The wild-type Bluetongue virus (BTV-WT), Bluetongue virus serotype 1 South African reference strain, GenBank accession numbers FJ969719 to FJ969728) was used for this study. The 10 genome segments were firstly cloned into pUC19 plasmid with T7 promoter and RNA transcripts sequentially prepared using the mMACHINE T7 transcription kit (Thermo). The three mutant viruses were generated by site-directed mutagenesis on BTV segment 8 or 9 as previously described, the start codons (ATG) of S8 were changed to GTG, preventing translation of NS2 for BTV-ΔNS2 (13), a calcium-binding site mutation (DDDE_250-253_AAAA) was introduced to prevent the formation of viral inclusion bodies (VIB) for BTV-ΔVIB (29), and the Entry-Competent-Replication-Abortive (BTV-ECRA) vaccine strain was designed with large deletions in an essential BTV gene encoding the VP6 protein (segment S9) of the internal core (16, 30). The antibodies produced in the laboratory were used for detection of viral protein NS2 (guinea pig) and VP6 (rabbit). Hoechst 33342 and Alexa 488-, 546-conjugated secondary antibodies were purchased from Life Technologies (Thermo).

### Reverse genetics, plaque assay and viral RNAs transfection

The wild-type and mutant viruses were recovered and reproduced (either from BSR, BSR-NS2 and BSR-VP6 cells) by reverse genetics as previously described (31). The plaque assay was undertaken with WT and mutant viruses to determine their titers. Viruses were diluted and applied to BSR, BSR-NS2, or BSR-VP6 cell monolayers at a multiplicity of infection (MOI) of 0.01 to 0.1, then overlaid with 1.2% Avicel. After 3 days, cells were fixed with formaldehyde, and plaque sizes were assessed. For colocalization analysis of BTV viral RNAs lacking both NS2 and VP6 in the host cytoplasm by HCR, BSR cells were transfected with all 10 BTV viral +ssRNA segments. The start codons (ATG) of segments S8 and S9 were mutated to GTG to prevent translation without significantly affecting nucleotide structure. Each +ssRNA segment (100 ng per well) was transfected into 12-well plates using X-TremeGENE HP DNA Transfection Reagent (Roche) according to the manufacturer’s instructions. Cells were incubated at 37□°C with 5% CO_₂_ for 20 hours prior to the HCR assay.

### Fluorescence *in-situ* Hybridisation Chain Reaction

RNA fluorescence *in-situ* Hybridisation Chain Reaction (HCR) was performed following established protocols (13). This two-step method employs initiator-tagged "Target" probes (30 oligonucleotides per RNA segment), designed using an online tool (https://www.biosearchtech.com), to bind and cover positive viral RNAs. An enzyme-free polymerization step then uses fluorescently labeled hairpin probes (H1 and H2) that form nicked double helices upon activation by the initiator strands (Supplementary Fig. 1). All oligonucleotide probes were synthesized by Integrated DNA Technologies, Inc. (see Supplementary Table 1). Cells were plated on poly-lysine-coated coverslips (VWR International) at a density of 2×10^5^ cells/well (12-well plate) and incubated overnight at 37 °C with 5% CO_₂_. Cells were then infected with BTV wild-type and mutant viruses at specified time points and MOIs. After infection, cells were washed with ice-cold PBSM (1× PBS, 5 mM MgCl_₂_), fixed with 4% paraformaldehyde in PBSM for 10 minutes at room temperature, and permeabilized with 0.1% Triton X-100 in PBSM for 10 minutes. After pre-incubation in pre-hybridisation buffer (2× SSC, 10% formamide) for 20 minutes, hybridisation buffer containing probes (5 ng/µL for the first step, 60 nM hairpin DNA for the second) was applied. The first hybridisation was carried out in a humidified chamber at 37 °C for 5 hours or overnight, followed by two washes with 10% formamide in 2× SSC at 37 °C.

Hairpin probes were annealed in a thermocycler (95 °C to 25 °C) before the second hybridisation, which was performed at 25 °C for 5 hours or overnight. After nuclear staining with 0.5 µg/mL Hoechst 33342, coverslips were mounted in ProLong Gold antifade media (Invitrogen) and examined by microscopy. For samples used for both immunofluorescence and HCR analyses, cells were first blocked with 1% BSA in PBSM for 1 hour at RT after fixation and permeabilization. The coverslips were then subjected to primary and secondary antibody staining in 1% BSA in PBS followed by another fixation step with 4% paraformaldehyde for 10 min. The cells were then washed once with PBSM and equilibrated with 10% formamide 2x SSC for 10 min before the *in-situ* hybridisation procedures. The entire experimental workflow is shown in the schematic overview illustrated in Supplementary Fig. 2.

### Image acquisition and analysis

Cells were imaged using 300 nm steps along the z-axis over a range of approximately 4.5 µm with a Nikon Ti2 wide-field microscope. The system was equipped with a 60× oil-immersion objective, 25 mm large wide-field-of-view (FOV) fluorescence capability, LED fluorescence and diascopic light sources, high signal-to-noise DAPI, GFP, and DS Red filters, as well as a high-sensitivity, large FOV monochrome CMOS camera. Quantification of the fluorescent photographs was performed at the same threshold and adjustment of contrast. Images were analysed in NIS-Elements.

### RNA complex pull-down assay

BSR cells were infected with BTV in MOI 5.0. The cells were lysed at 5 h post infection with RNA complex buffer (20mM Tris-HCl pH 7.5, 200mM KCl, 5mM magnesium acetate, 1mM DTT, Protease inhibitor (Merck), RNase inhibitor (Thermo), 0.1% Triton X-100 (32) and the nucleoli removed by centrifugation. The supernatant was mixed with streptavidin agarose beads (Novagen) coated with a biotin-labelled primers which annealed the coding region of BTV-1 S1, S6, or S10 (supplementary table 2). The beads were incubated with poly-A RNA first, to decrease non-specific binding. After 30 min incubation at 28°C, the beads were washed three times with excess buffer followed by 1 min heating at 80°C to release the RNA. The eluted RNA samples were then subjected to qRT-PCR to measure the quantities of ten segments from each kind of beads. Genomic dsRNA extracted from BTV was used as the equal molar ratio standard for qRT-PCR. cDNA synthesis was performed using the SuperScript™ III Reverse Transcriptase Kit (Invitrogen), following the manufacturer’s instructions. qPCR was carried out with PowerUp™ SYBR™ Green Master Mix (Thermo Fisher), and the primers used are listed in Supplementary Table 3.

## RESULTS

### Identification and visualization of intersegments interaction and RNA complex formation in BTV infected cells

BTV RNA segments interact with each other *in vitro* and form RNA complexes that are important for RNA packaging. To investigate, whether such RNA complexes are formed during virus replication in the infected cells, we used a highly efficient RNA fluorescence *in-situ* Hybridisation Chain Reaction (HCR) technique tailored for BTV RNAs (13). Following hybridisation of specific oligonucleotide probes to the +ssRNA segments of the virus, an enzyme-free polymerization reaction is used to amplify a strong fluorescent signal (Supplementary Fig. 1). HCR analysis was performed on BSR cells infected with wild type BTV (BTV-WT) at a multiplicity of infection (MOI) of 100 and analysed 8 hours post-infection (hpi). Two probe sets, labelled with Cy3 (green, 550 nm) and Cy5 (red, 670 nm), targeting the BTV RNA segments S6 and S10, were employed (Fig. 1A & Supplementary Fig.1B). Since S7-S10, the four smaller RNA segments form complexes first (15), which then recruit S6, S10 and S6 probes were used to for the identification of the RNA complex formation. Quantitative measurements revealed that over 80% of Cy3 and Cy5 spots colocalized within the cells (Fig. 1B) indicative of complex formation in infected cells. Two control experiments were undertaken to validate the results further. For the positive control, two probe sets targeting different regions of the same RNA segment (S10) and labelled with Cy3 and Cy5 were used. In BSR cells infected with BTV-WT and analysed at 8 hpi, S10 RNA was detected by both probes (Fig. 1A). As expected, quantitative results showed over 90% colocalization of Cy3 and Cy5 spots (Fig. 1B), supporting the specificity of the analysis. To ensure that colocalization efficiency was not influenced by high concentration of viral RNA, a negative control was performed using labeled probes targeting the highly expressed cellular β-actin mRNA (ActB) and viral RNA S10. In BSR cells infected with BTV-WT and hybridised with Cy3-labeled probes for ActB and Cy5-labeled probes for S10 at 8 hpi, minimal colocalization of these RNA species was observed (Fig. 1A). Quantitative analysis confirmed a low colocalization percentage of ≤22% (Fig. 1B). These results validate the specificity and sensitivity of the HCR technique and establish robust positive and negative controls for the quantitative analysis.

**Fig 1.**
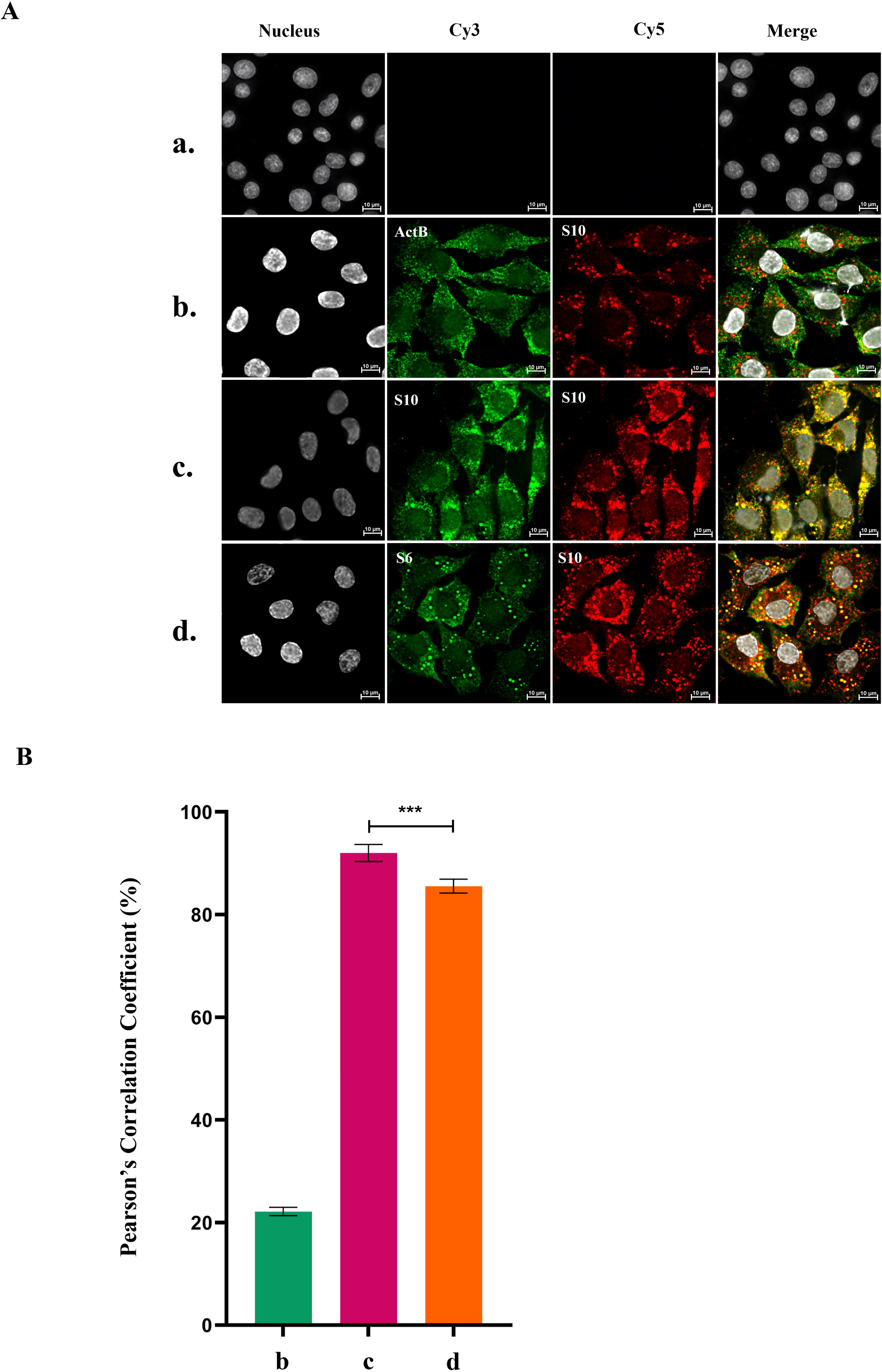
*In-situ* Hybridisation Chain Reaction (HCR) and colocalization analysis of Bluetongue virus (BTV) in BSR cells. **(A)** BSR cells were infected with wild-type Bluetongue virus (BTV-WT) at a multiplicity of infection (MOI) of 100 for 8 hours in 12-well plates, then fixed and incubated with probes targeting cellular mRNA or BTV +ssRNA segments for HCR and colocalization analysis. Uninfected BSR cells served as the mock control (row a). For the negative control (row b), cells were incubated with probes targeting cellular mRNA β-actin (ActB, green, Cy3, 550 nm) and BTV +ssRNA segment S10 (red, Cy5, 670 nm). The positive control (row c) utilized Cy3-and Cy5-labeled probes targeting different regions of segment S10 to validate colocalization and HCR. For row d, probes labelled with Cy3 and Cy5 were used to target BTV RNA segments S6 and S10. Hoechst 33342 was used to stain cell nuclei. Colocalization analysis revealed yellow spots, indicating the overlap of Cy3 and Cy5 signals. **(B)** Pearson’s correlation coefficients were quantified to assess colocalization between Cy3 and Cy5 spots for each experiment. Analysis was conducted across five images from separate replicates, representing over 100 cells per experiment. The error bars represent standard deviations. ***: t-test p value < 0.005.

To substantiate that the RNA segments indeed form RNA complexes in the infected cells, we designed a biotinylated oligos-based pull-down assay, similar to the one previously used for *in vitro* study (15). At 5 hours post-infection, when viral proteins were synthesised at low level only, the RNA-RNA interactions and RNA complex formation were detected by pulled-down beads assay using three representative RNA segments (S1, S6, or S10), as described in Methods. The qRT-PCR data clearly identified BTV RNA complexes (Fig 2). The beads-coated with each of the three RNA segments pulled down remaining nine RNA segments almost at equal efficiency, indicating that complexes containing all segments exist in early stage of virus replication.

**Fig 2.**
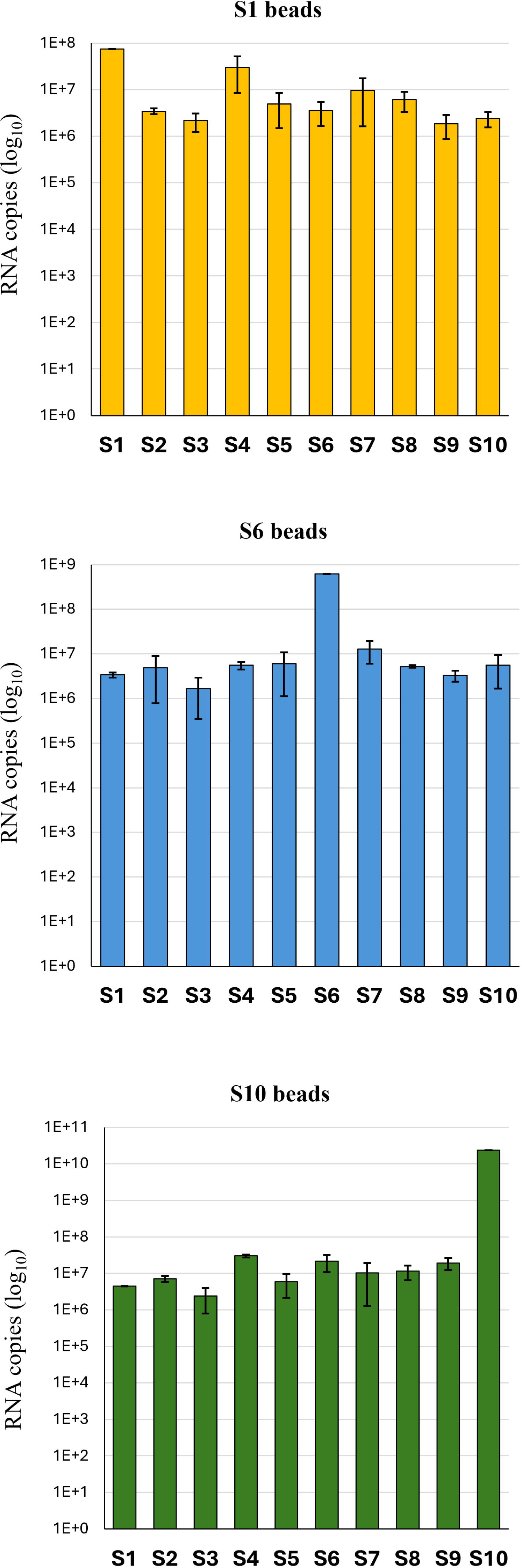
RNA complexes containing all segments could be pulled down from BTV infected cell lysate. BSR cells were infected with BTV-1 at MOI 5.0 for 5 hours. Beads coated with a S1-, S6-, or S10- specific oligos were incubated with the cell lysate for 30 mins to interact with BTV RNAs in the lysate; bound RNAs was then sequentially extracted. The amount of each interacting RNA segment was determined by qRT-PCR using primers specific to the indicated segment, correlated to an equal molar ratio BTV genomic RNA standard. The RNA copy number per ml of infected call lysate was shown in log_10_. Standard deviation from multiple individual experiments were calculated and shown as error bars.

### NS2 is not essential for BTV RNA complex formation although it facilitates colocalization of larger RNA segments in infected cells

To investigate the correlation between NS2 and RNA packaging in infected cells, two mutant BTV viruses were generated through site-directed mutagenesis of segment 8, the NS2 encoding BTV RNA segment, as previously described. The start codons (ATG) of segment 8 were altered to GTG, preventing the translation of NS2 in BTV- ΔNS2 (13). Additionally, a calcium-binding site mutation (DDDE_250-253_AAAA) was introduced to block the formation of viral inclusion bodies (VIB) in BTV-ΔVIB (29). Rescue of the two mutant strains BTV-ΔVIB and BTV-ΔNS2, was only successful in a NS2-complementing cell line (BSR-NS2) (Fig. 3A). To investigate the colocalization kinetics of different viral RNA species during the BTV replication, we visualised S10 and S6, two of the earliest RNAs to be recruited but also, importantly, segment S1 which our previous *in vitro* studies have shown is the last segment to be incorporated, marking completion of the set of segments comprising the genome. We performed HCR and colocalization analysis on three viral +ssRNA segments: S1 (large-sized), S6 (medium-sized), and S10 (small-sized), in the wild-type (BTV-WT) and two mutant strains (BTV-ΔVIB and BTV-ΔNS2) at 20 hpi with a MOI 5.0. Viral RNA segments were detected using HCR with S1 labeled in blue (Atto-488, 499 nm), S6 in green (Cy3, 550 nm), and S10 in red (Cy5, 670 nm). Colocalization analysis revealed yellow spots, indicating overlap between Cy3 and Cy5 signals, while pink, purple, or white spots represented colocalization among Atto-488, Cy3, and Cy5 signals (Fig. 3B). Pearson’s correlation coefficients were calculated to quantify colocalization among viral RNAs S1, S6, and S10 in the cells infected with WT and two mutant BTV strains. The results revealed the highest colocalization efficiency between S6 and S10 for all strains, with Pearson’s correlation coefficients ∼90%, with WT showing slightly a higher colocalization efficiency compared to BTV-ΔVIB and BTV-ΔNS2 (Fig. 3C). BTV-ΔNS2 still exhibited colocalization although affected the efficiency between S1 and S10, as well as S1 and S6, with Pearson’s correlation coefficients 65%. However, BTV-ΔVIB (but still retaining NS2) showed significantly higher colocalization efficiency between these RNA pairs (Fig. 3C). These findings suggest that while VIBs and NS2 are essential for BTV replication in BSR cells (9), viral RNA segments can still form complexes in the cytoplasm independently of these structures. These findings of *in vivo* infected cells are consistent with the previous *in vitro* studies (13, 15) and also indicate that while NS2 is not required for assembling smaller RNA segments, it may influence the assembly of larger BTV RNA segments.

**Fig 3.**
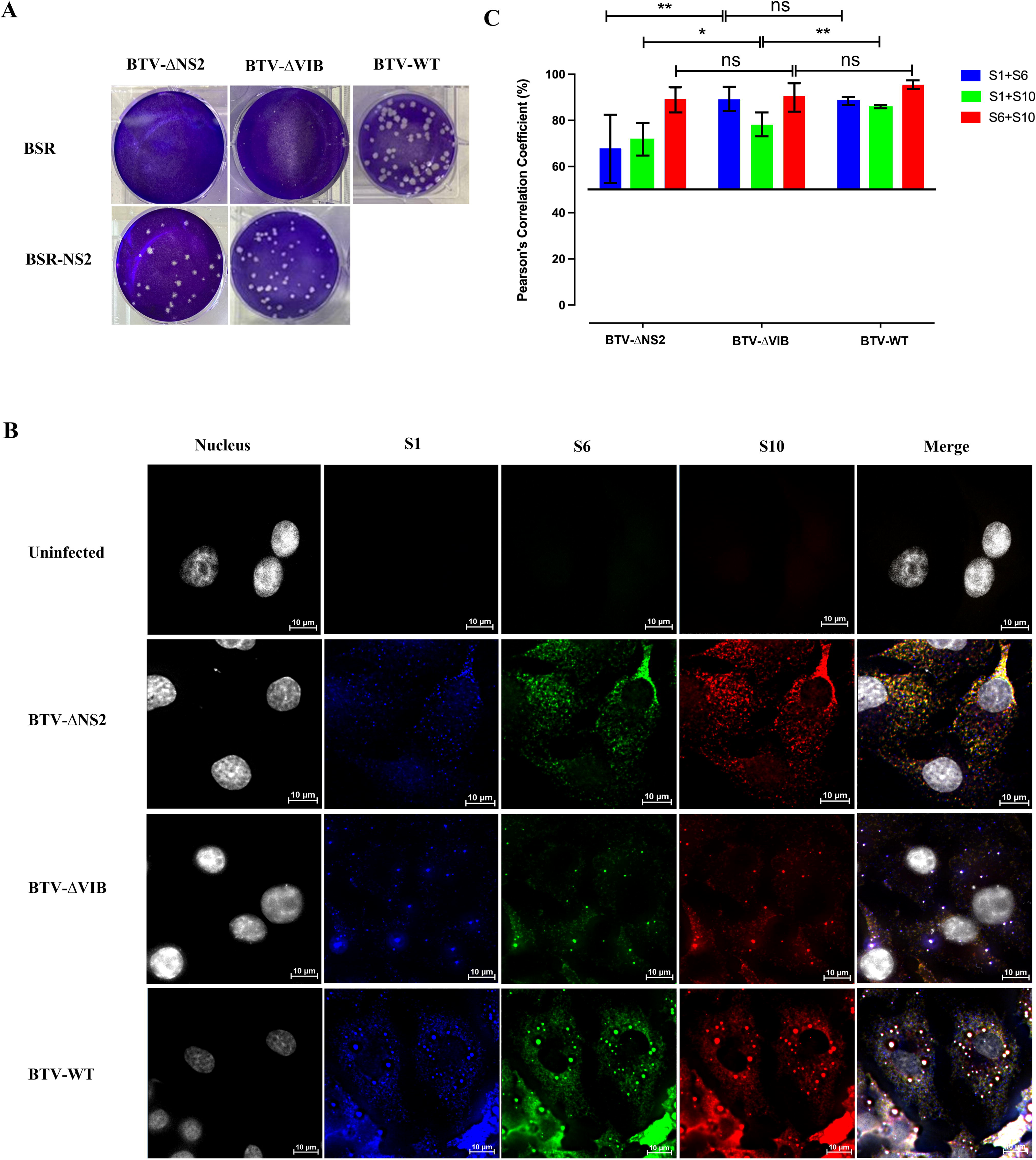
Colocalization analysis of viral RNAs in BSR cells infected with wild-type and mutant BTV using HCR. **(A)** Plaque assay results for BTV-WT and two mutant strains (BTV-ΔVIB and BTV-ΔNS2) in BSR and BSR-NS2 cell lines. **(B)** BSR cells were infected with BTV-WT, BTV-ΔVIB and BTV-ΔNS2 at a MOI 5.0 for 20 hours in 12-well plates. Uninfected BSR cells served as a negative control. Viral RNA segments S1 (blue, Atto-488, 499 nm), S6 (green, Cy3, 550 nm), and S10 (red, Cy5, 670 nm) were detected using HCR. Colocalization analysis showed yellow spots indicating overlap between Cy3 and Cy5 signals, while pink, purple, or white spots represented colocalization of Atto-488, Cy3, and Cy5 signals. Hoechst 33342 was used to stain cell nuclei. **(C)** Quantification of Pearson’s correlation coefficients for colocalization between viral RNAs S1, S6, and S10. Six images from independent replicates, covering over 120 cells per experiment, were analysed. The error bars represent standard deviations. ns: un-paired t-test no significance. *: t-test p value < 0.05. **: t-test p value < 0.01.

### Smaller segments of BTV viral RNAs form complexes prior to the larger segments

To investigate the sequence of RNA complex network formation during BTV replication, BSR cells were seeded in 12-well plates and infected with either BTV-WT (Fig. 4A) at an MOI of 100 or the mutant strain BTV-ΔNS2 (Fig. 4C) at an MOI of 5.0. Cells were fixed at various time points post-infection for HCR and colocalization analysis. BTV S1 (blue, Atto-488, 499 nm), S6 (green, Cy3, 550 nm), and S10 (red, Cy5, 670 nm) were detected using HCR. Colocalization analysis revealed yellow spots (overlapping Cy3 and Cy5 signals), while pink, purple, or white spots indicated colocalization of Atto-488, Cy3, and Cy5 signals. The number of colocalized spots for all viral RNA segments increased over time in both BTV-WT and BTV-ΔNS2 infected BSR cells (Fig. 4A and C). Pearson’s correlation coefficients remained consistently higher for the smaller viral RNA segments, S6 and S10, across nearly all post-infection time points in both BTV-WT and BTV-ΔNS2 infected cells albeit with a decrease at 7 hpi and 8 hpi in BTV-WT infected cells, due to segment entry into assembled VIBs (Fig. 4B and D). Additionally, BTV-ΔNS2 exhibited lower complex formation efficiency, with significantly lower Pearson’s correlation coefficients for larger segments compared to BTV-WT (Fig. 4B and D), confirming that smaller BTV RNA segments assemble into complexes before larger ones. These findings highlight the critical role of NS2 in facilitating the assembly of larger RNA segments.

**Fig 4.**
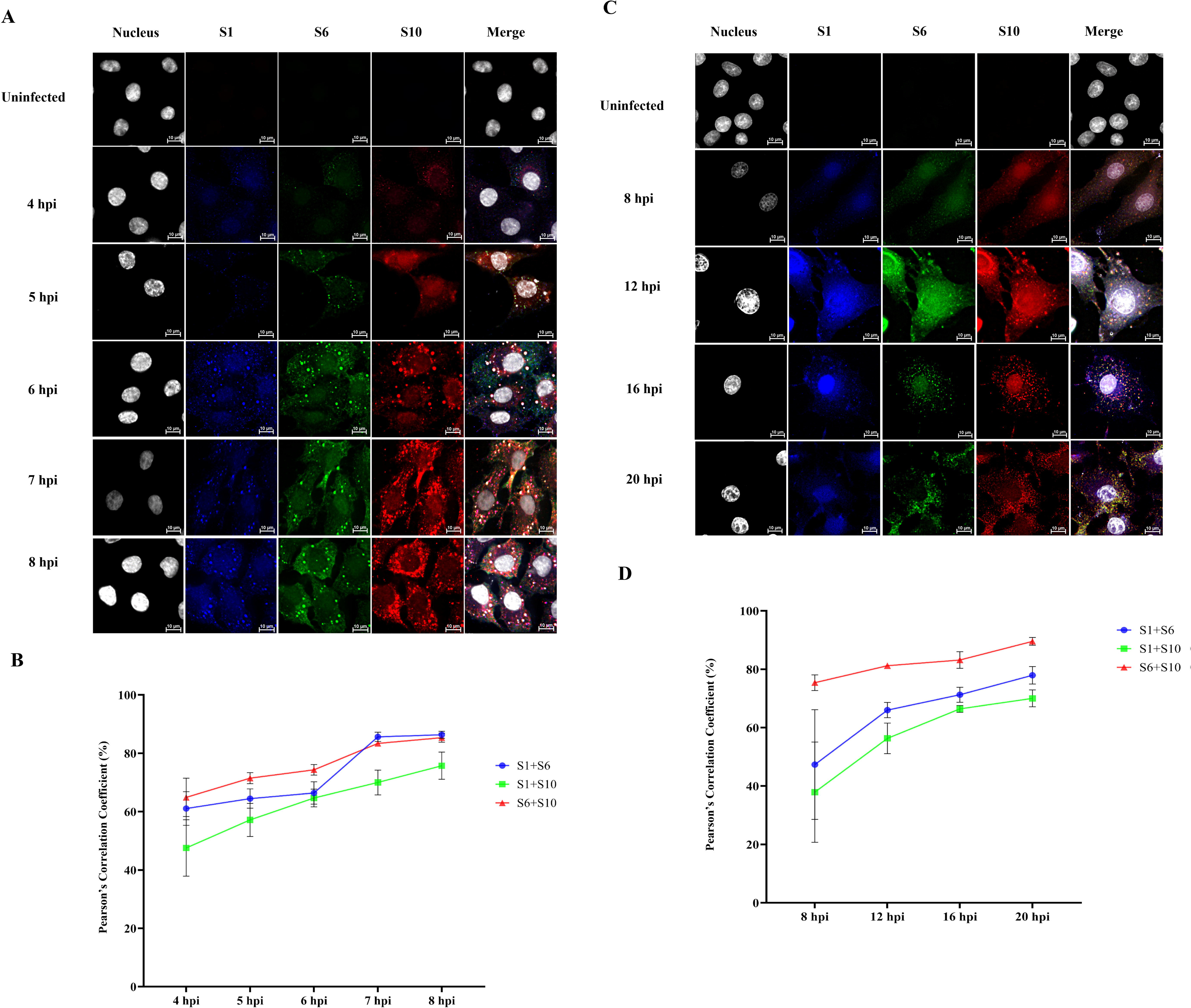
Colocalization analysis of viral RNAs in BSR cells infected with BTV-WT and BTV-ΔNS2 at different post-infection time points. BSR cells in 12-well plates were infected with BTV-WT **(A, B)** at a multiplicity of infection (MOI) of 100 or with BTV-ΔNS2 **(C, D)** at a MOI of 5.0, and fixed at various time points post-infection for HCR and colocalization analysis. Uninfected BSR cells served as a negative control. Viral RNA segments S1 (blue, Atto-488, 499 nm), S6 (green, Cy3, 550 nm), and S10 (red, Cy5, 670 nm) were detected using HCR. **Panel A** shows representative HCR images of BTV-WT-infected cells, revealing colocalization of viral +ssRNA segments as yellow (Cy3 + Cy5), pink/purple, or white spots, indicating partial or complete overlap of signals. Corresponding quantitative analysis in **panel B** presents Pearson’s correlation coefficients for S1, S6, and S10 RNA colocalization across time points in BTV-WT-infected cells. **Panel C** shows the same HCR analysis as in panel A, performed on BTV-ΔNS2-infected cells at a lower MOI, with similar visualization of RNA colocalization. **Panel D** provides the quantitative Pearson’s correlation analysis for BTV-ΔNS2 infection, as shown in panel C. For all conditions, five images per sample from independent replicates, covering over 100 cells per experiment, were analysed. Error bars represent standard deviations.

### VP6 may play a key role in BTV viral RNAs assembly prior to recruitment into VIBs

VP6 has been identified as a crucial component of the primary replicase complex and is hypothesized to play a role in recruiting and packaging BTV RNA (9, 10). To confirm the correlation between VP6 and RNA packaging, BSR cells were infected either with BTV-WT, or BTV-ΔVIB or BTV-ΔNS2 at a multiplicity of infection (MOI) of 5.0 for 20 hours in 12-well plates and colocalizations of RNAs and viral proteins were examined using HCR and immunofluorescence. Viral +ssRNA segments S10, S6, and S1, as well as cellular mRNA ActB, were detected using HCR (red, Cy5, 670 nm). Immunofluorescence localized the viral structural protein VP6 (green, Alexa-546, 561 nm) and NS2 (blue, Atto-488, 499 nm). Yellow spots indicate colocalization between Alexa-546 and Cy5 signals, while pink, purple, or white spots represent colocalization between Atto-488, Alexa-546, and Cy5. The results showed that the VP6 protein colocalized with BTV viral +ssRNA segments S1, S6, and S10 of the RNA complex, as well as with cellular mRNA ActB, in BSR cells infected with BTV-WT, BTV-ΔVIB, or BTV-ΔNS2 (Fig. 5). This confirms VP6 as a general RNA binding protein consistent with previous *in vitro* studies (9, 17, 33). Notably, VP6 colocalized with both viral +ssRNA and VIBs in BTV-WT infected cells and with defective VIBs in BTV-ΔVIB-infected cells but did not colocalize with non-aggregated NS2 protein (Fig. 5A and B). Additionally, VP6 colocalized with viral +ssRNA and VIBs but was absent from VIBs associated with cellular mRNAs (Fig. 5). These findings suggest that VP6 binds viral and cellular RNAs independently of NS2 and that its colocalization with VIBs likely reflects binding to the viral +ssRNA accrued there. To confirm this, BSR cells were infected with the BTV-ECRA mutant strain (16), which has large deletions in segment S9 preventing VP6 translation, at an MOI of 5.0 for 20 hours. Colocalization analysis of viral RNA segments S1, S6, and S10 (detected using HCR) demonstrated that viral RNA complexes still formed and entered VIBs despite the absence of VP6 (Fig. 6). These findings indicate that VP6 can associate with viral +ssRNA before they enter VIBs, and that it may play a role in RNA recruitment and early assembly although it is not essential for RNA entry into VIBs. This suggests that BTV RNA packaging into VIBs can occur through alternative mechanisms, emphasizing the complexity of the BTV replication process.

**Fig 5.**
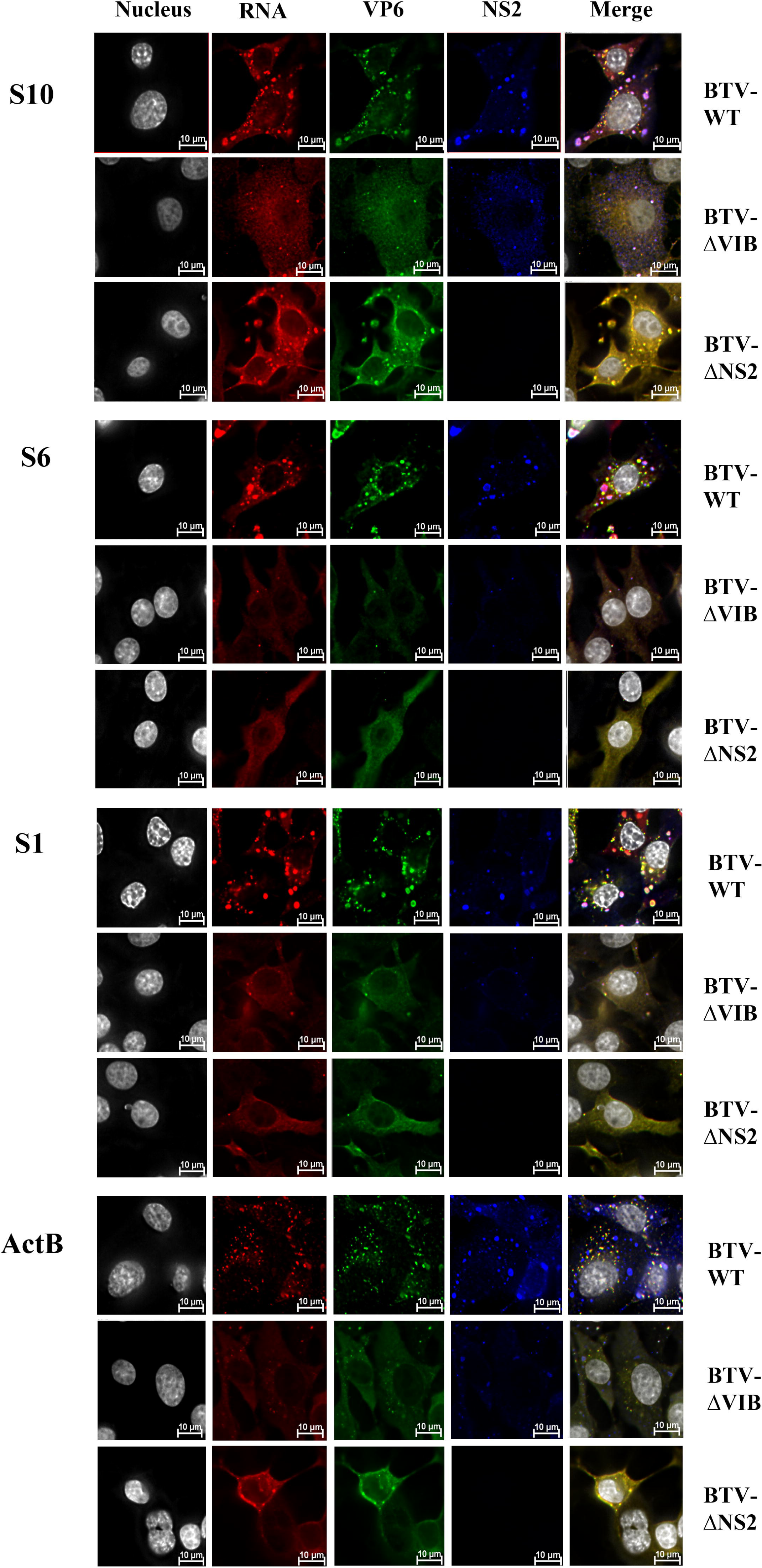
BTV structure protein VP6 colocalized with viral and host RNAs. BSR cells were infected with BTV-WT, BTV-ΔVIB and BTV-ΔNS2 at a MOI 5.0 for 20 hours in 12-well plates. Uninfected BSR cells served as a negative control. Cell nuclei were stained with Hoechst 33342. Colocalization of RNAs and viral proteins was examined using HCR and immunofluorescence. Viral +ssRNA segments S10, S6, and S1, as well as cellular mRNA β-actin (ActB), were detected using HCR (red, Cy5, 670 nm). Immunofluorescence localized the viral structural protein VP6 (green, Alexa-546, 561 nm) and the non-structural protein NS2 (blue, Atto-488, 499 nm). Yellow spots indicate colocalization between Alexa-546 and Cy5 signals, while pink, purple, or white spots represent colocalization between Atto-488, Alexa-546, and Cy5. Five images from separate replicates, encompassing over 100 cells per experiment, were analysed.

**Fig 6.**
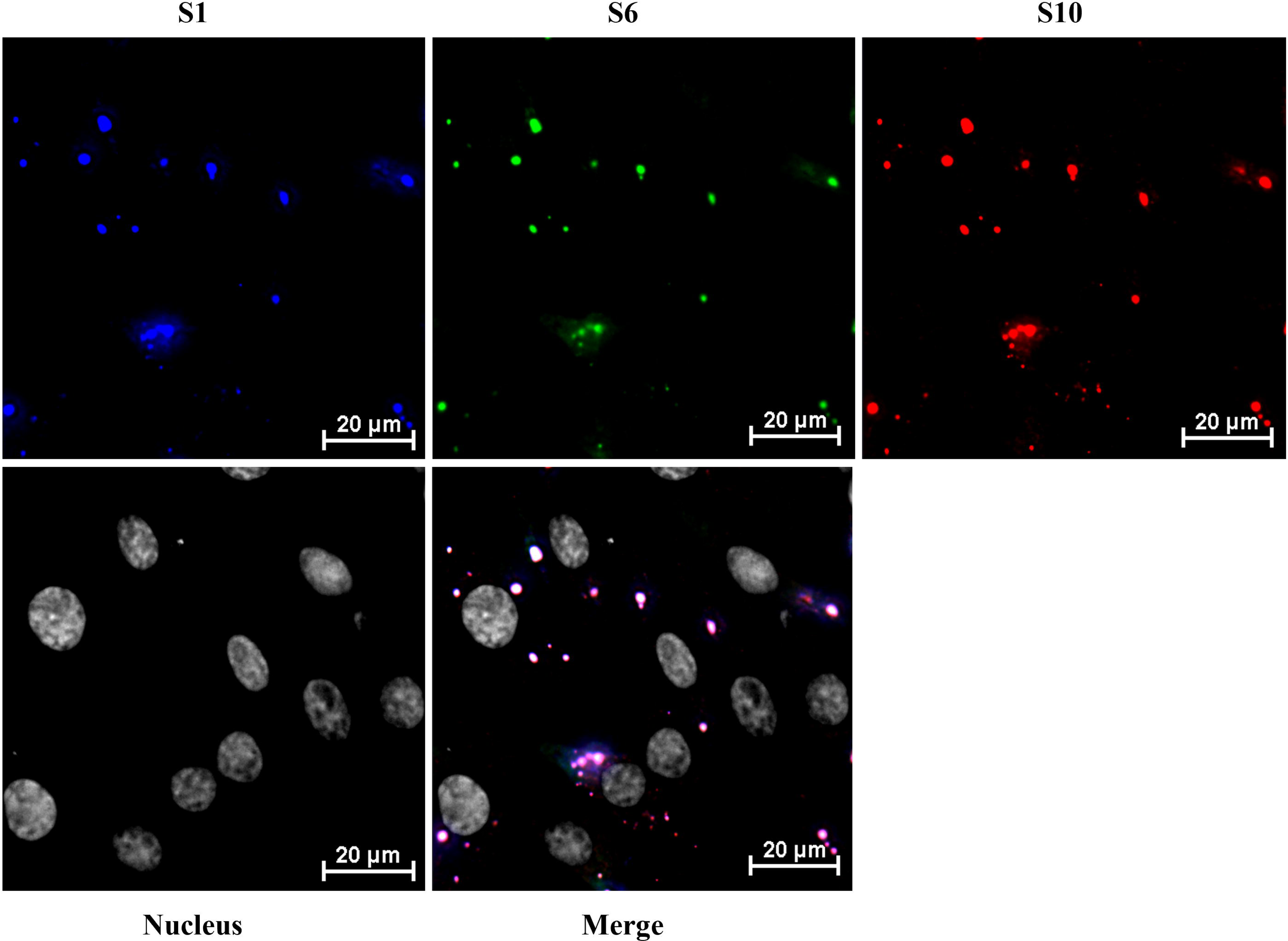
BTV viral RNAs enter VIBs independently of VP6. BSR cells were infected with mutant strains (BTV-ECRA) at a MOI 5.0 for 20 hours in 12-well plates. Viral RNA segments S1 (blue, Atto-488, 499 nm), S6 (green, Cy3, 550 nm), and S10 (red, Cy5, 670 nm) were detected using HCR. Colocalization analysis showed yellow spots indicating overlap between Cy3 and Cy5 signals, while pink, purple, or white spots represented colocalization of Atto-488, Cy3, and Cy5 signals. Hoechst 33342 was used to stain cell nuclei. Five images from individual replicates, encompassing over 100 cells per experiment, were analysed.

### Complete set of RNA genome selection for packaging do not require either VP6 or NS2

To investigate whether BTV +ssRNA segments can form RNA complexes in the host cell cytoplasm in the absence of both NS2 and VP6, BSR cells were transfected with all 10 BTV viral +ssRNA segments. The start codons (ATG) of segments S8 and S9 were mutated to GTG, preventing translation without significantly altering nucleotide structure. Using HCR, RNA segments S1 (blue, Atto-488, 499 nm), S6 (green, Cy3, 550 nm), and S10 (red, Cy5, 670 nm) were detected in the cytoplasm. Despite the absence of NS2 and VP6, RNA complexes still formed in the cytoplasm. Pearson’s correlation coefficients remained high (∼80%) for the smaller RNA segments S6 and S10, while larger segments, such as S1, showed lower correlations (∼60%), mirroring the patterns observed in BSR cells infected with BTV-ΔNS2 (Fig. 7). These results, consistent with previous *in vivo* assays (infected cells), further support the notion that RNA packaging into a complex is an innate feature of the sequence but that it may be enhanced by the virally encoded RNA binding proteins.

**Fig 7.**
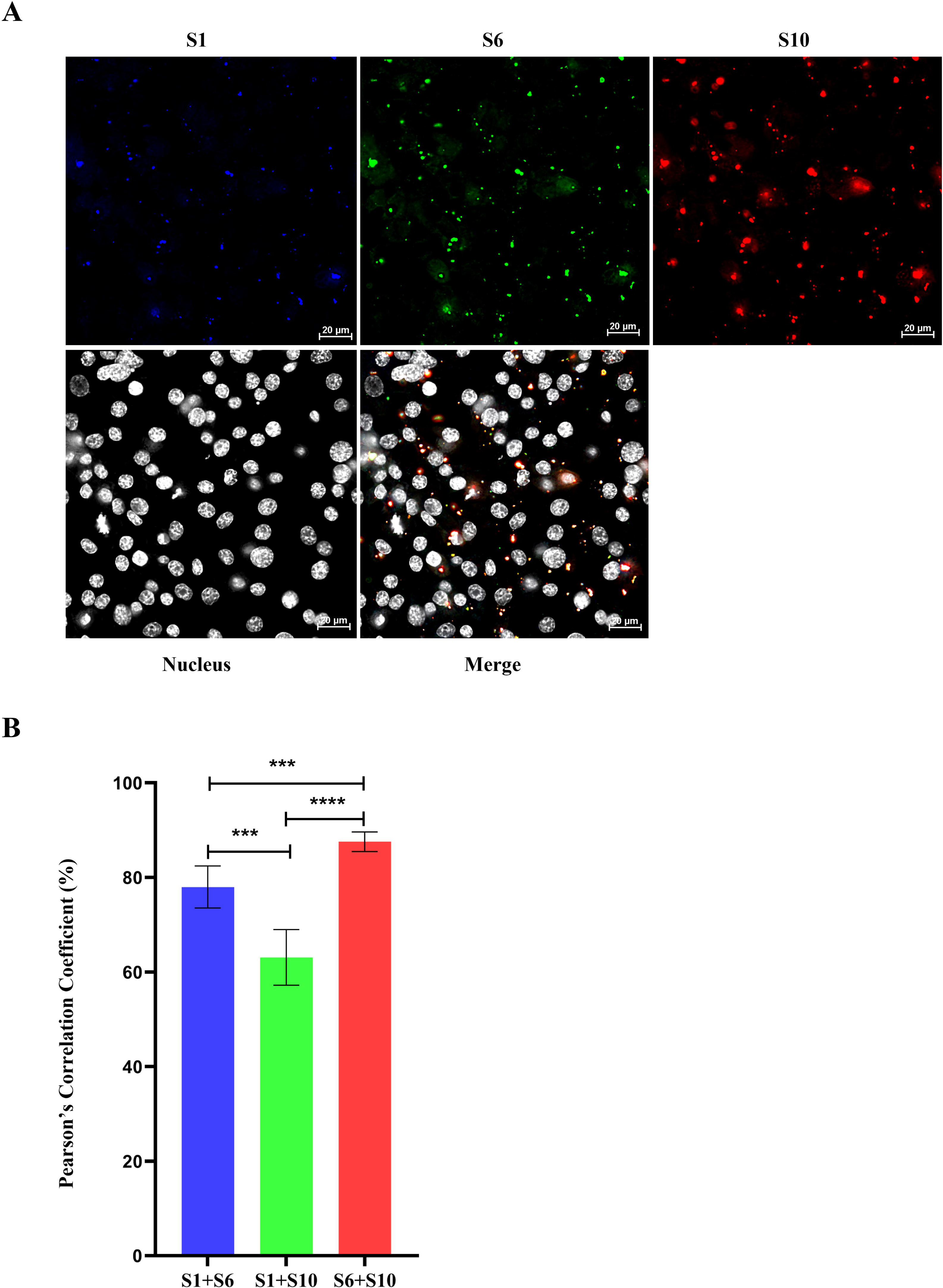
Colocalization analysis of BTV viral RNAs in the absence of both NS2 and VP6 in the host cell cytoplasm by HCR. BSR cells were transfected with all 10 BTV viral +ssRNA segments. The start codons (ATG) of segments S8 and S9 were mutated to GTG, preventing translation without significantly altering nucleotide structure. **(A)** Viral RNA segments S1 (blue, Atto-488, 499 nm), S6 (green, Cy3, 550 nm), and S10 (red, Cy5, 670 nm) were detected by HCR in the cytoplasm. **(B)** Quantification of Pearson’s correlation coefficients for colocalization between viral RNAs S1, S6, and S10. Five images from individual replicates containing more than 200 cells were analysed for each experiment. The mean standard error bars are shown. ***: t-test p value < 0.005, ****: t-test p value < 0.001.

## DISCUSSION

The selection of viral genomes from the pool of cellular nucleic acids and their subsequent packaging within the virus capsid are two critical processes in virus infected host cells. The findings presented in this study provide, for the first time, how multipartite BTV genomic RNAs are assorted in the infected cells by RNA-RNA interactions to form specific RNA network. Utilizing the HCR technique for *in-situ* hybridisation and colocalization analysis, it was possible to show the dynamic nature of BTV +ssRNAs interactions within infected BSR cells. Our results from BTV infected cells confirm the viral RNA segments indeed interact with each other to form the RNA complex of ten segments as previously observed in our *in vitro* studies (13, 15).

Our current *in vivo* findings are consistent with our earlier *in vitro* studies, which support a model where BTV RNA packaging is orchestrated through the stepwise formation of a complex RNA-RNA interaction network. This process appears to follow a defined sequence, beginning with the smallest segment, S10, which initiates interactions with other small segments to form an initial complex (S7–S10). Medium-sized segments (S4–S6) and eventually the larger segments (S1–S3) are then recruited in a hierarchical manner, leading to the complete genome complex ready for encapsulation. However, the observed association patterns *in vivo*, such as S6 colocalizing with both S1 and S10 at similar level, while S1 and S10 show weaker direct association, suggest that the assembly pathway may be more dynamic or flexible within the cellular environment than what is captured in simplified *in vitro* systems. It is possible that S6 functions as a central scaffold or bridging segment, facilitating transient or stage-specific interactions between small and large segments, even if those segments are not simultaneously colocalised. These findings raise the possibility of alternative or parallel assembly routes and intermediate complexes that may form depending on spatial constraints or temporal regulation within the cytoplasm. Exploring whether larger segments interact directly or form subcomplexes before engaging with the initial S7–S10 complex remains an open question and will be an important focus of future investigations.

Further, in BSR cells infected with the NS2-deficient mutant BTV-ΔNS2, we observed RNA segments could still form complexes in the absence of NS2, although there was a reduction in the colocalization efficiency of viral RNA segments, especially between large and small RNA segments. This finding aligns with previous reports demonstrating that NS2 facilitates the formation of viral inclusion bodies (VIBs), which serve as scaffolds for RNA recruitment (29, 34). Thus, although VIBs and NS2 are not directly involved in RNA-RNA interaction and complex formation, they are key components for the efficient assembly of the viral genome, particularly for larger RNA segments, despite being dispensable for the initiation of RNA complex formation.

Our study provides new evidence supporting a key role for VP6 in BTV RNA packaging and assembly. VP6, a structural protein previously identified as a core component of the BTV replicase complex (17, 33), was found to colocalize with both viral and cellular RNAs independently of NS2 and VIBs in cells infected with mutant strains (BTV-ΔVIB and BTV-ΔNS2), and also to associate with VIBs in wild-type BTV (BTV-WT) infection. This pattern suggests that VP6 associates with RNA network prior to the formation of VIBs. Interestingly, VP6 was only recruited to VIBs in the presence of viral RNAs but not cellular RNAs, implying a selective mechanism that favours viral RNAs. Furthermore, VP6 binds viral RNAs early in infection, indicating its involvement in the initial stages of RNA packaging and suggesting it may act as a checkpoint to regulate RNA complex formation and prevent potential cytotoxicity. These findings, consistent with previous *in vitro* studies (10, 17, 33), highlight the multifunctional role of VP6 in the BTV replication cycle.

Direct confirmation of RNA-RNA interactions and RNA network formation do not require NS2 or VP6 was further demonstrated in cells transfected with all ten BTV +ssRNA segments, where RNA complexes still formed with high efficiency (∼80%) for smaller segments (S6-S10), while larger segments such as S1 showed less efficiency (∼60%), similar to results observed in BTV-ΔNS2 infections. These findings suggest that BTV RNA segment interactions are driven by intrinsic RNA-RNA affinities, with smaller segments initiating complex formation followed by the incorporation of larger segments (e.g., S1). This supports the notion of a sequential RNA-RNA interactions and packaging model, consistent with earlier *in vitro* studies (13, 15). The use of the HCR technique enabled precise localization and quantification of these interactions, suggesting that RNA-RNA interactions and RNA complex formation facilitate the initial stages of genome packaging. Notably, results obtained from in *vivo* infected cells indicate that while RNA complex formation occur independently of NS2, the presence of NS2 enhances assembly efficiency, particularly, of the larger RNA segments suggesting that NS2 might play an important role in concentrating RNAs and orchestrating the formation of the complete genome structure during virus replication.

Altogether, our findings support a model in which BTV RNA segments are assorted via highly specific intersegments interactions while VP6 associates with the interacting RNA segments and RNA complexes. The VP6-RNA complexes are recruited to VIBs via NS2, the major component of VIBs, within which core assembly and RNA packaging are orchestrated by RNA-protein and protein-protein interactions. VP6 may play a role in directing these RNAs into viral inclusion bodies (VIBs). NS2, in turn, functions as a scaffolding protein within VIBs, organizing and concentrating viral RNAs and proteins to facilitate core assembly. Both VP6 and NS2 appear to enhance and stabilize the packaging process rather than initiate it and may also serve a protective function probably necessary for shielding viral RNAs from degradation.

In summary, our findings reveal that BTV RNA packaging is governed by a complex, stepwise mechanism. While smaller RNA segments appear to initiate complex formation early, NS2 facilitates the recruitment and coordination of larger segments within the assembly factories, the VIBs and VP6 binds viral +ssRNAs early in infection, potentially guiding them into VIBs for packaging into the assembling inner capsid (VP1, VP3, VP4, VP6) and subsequent core formation. These findings are highly significant for BTV but also offer broader implications for the packaging strategies of other dsRNA viruses with segmented genome. Despite these advances, our study has limitations. This study marks the first application of HCR in dsRNA virus research, where it demonstrated high sensitivity and specificity in detecting BTV RNAs, both in transfected and infected BSR cells. However, its application is currently limited to fixed-cell imaging, restricting its use for real-time RNA tracking. Although live-cell imaging approaches have been developed in studies of other viruses to provide dynamic insights into RNA interactions (35), their application in BTV and related viruses has been limited to protein tracking in live infected cells (36), with RNA tracking techniques remaining largely underdeveloped. Additionally, colocalization analysis alone does not establish direct evidence of interaction. Probe accessibility may also influence RNA detection efficiency, and validation in primary cells or animal models is needed to confirm our findings. Nevertheless, HCR is a powerful tool for investigating viral RNA interactions in infected cells. Future research should further dissect the molecular interplay between viral proteins and RNA segments, particularly the precise role of VP6 in RNA recruitment, to better understand.

## AUTHOR CONTRIBUTIONS

D.-S.L. and P.R. designed the experiments. D.-S.L. and P.-Y.S. conducted the experiments and analysed the data. D.-S.L. and P.R. drafted the manuscript, while D.-S.L., P.-Y.S. and P.R. contributed to its review and editing. P.R. provided supervision, managed the project, and secured funding.

## ACKNOWLEDGMENTS

We gratefully acknowledge Dr. Zhi-Jian Zhou and Prof. Xing-Yi Ge from Hunan University, Changsha, China, for their helpful suggestions and support with data analysis during the early stages of the project.

This project was supported by a Wellcome Trust Senior Investigator Award (221749/Z/20/Z) awarded to P.R.

## SUPPLEMENTARY DATA

Supplementary Data are available online.

## DATA AVAILABILITY

Data available upon request.

