## Supplemental figures and tables for "Visualization of Bluetongue virus RNA segment networks in infected cells: multipartite genomic RNA assortment is independent of viral proteins NS2 and VP6"

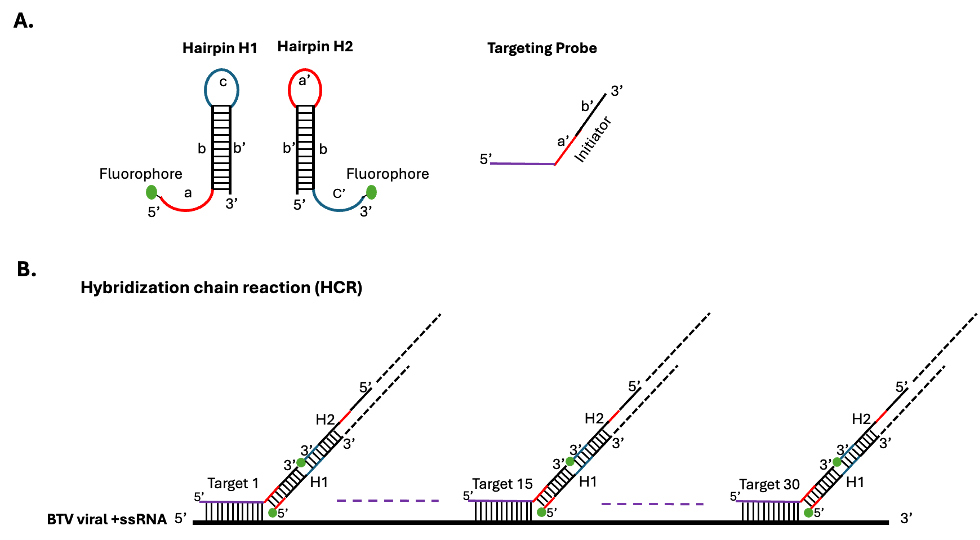

**Supplementary Fig 1. The cartoon shows the *in-situ* Hybridisation Chain Reaction (HCR), previously published (**Sung et al., 2024**). (A)**The structures of H1 and H2 of hairpin DNA pairs, and corresponding targeting probe including initiator sequence. **(B)** The *in-situ* Hybridisation Chain Reaction (HCR) strategy: the 30 targeting probes spanning the BTV viral +ssRNA segment, each including an initiator sequence hybridise to DNA hairpin H1 and then H1 and H2 hairpins continue to hybridise with each other.

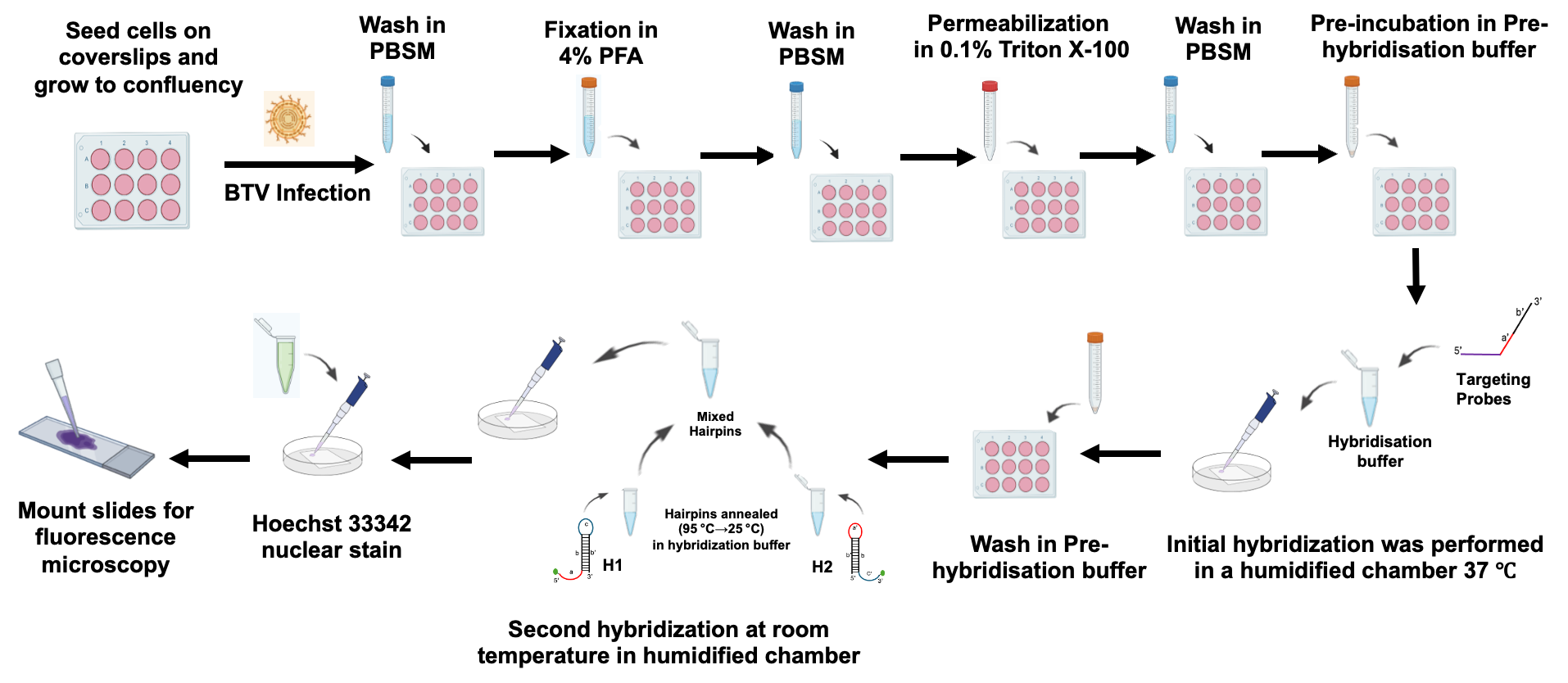

**Supplementary Fig 2. The illustration depicts a schematic overview of the *in-situ* hybridisation chain reaction (HCR) experiments conducted.**

**Supplementary Table 1. RNA probes used for *in-situ* Hybridisation Chain Reaction**

| S1-HCR-probes-atto488 | Probe (5'-> 3') | Probe position | Percent GC |
| --- | --- | --- | --- |
| RNA-HCR-Probe-H1_5'-Atto488 | Atto488 - 5' - TAGACTGAACCCACTCCGACG***ATCTGTCTT***CGTCGGAGTGGG - 3' |  |  |
| RNA-HCR-Probe-H2_3'-Atto488 | 5' - CGTCGGAGTGGG***TTCAGTCTA***CCCACTCCGACGAAGACAGAT - 3' - Atto488 |  |  |
| BTV1-WT-S1_RNA-HCR-target_1 | aacctcatctaatactggtatc***AAA***CGTCGGAGTGGGTTCAGTCTA | 227 | 36.00% |
| BTV1-WT-S1_RNA-HCR-target_2 | cctcaaaagcgtaactttggaa***AAA***CGTCGGAGTGGGTTCAGTCTA | 279 | 41.00% |
| BTV1-WT-S1_RNA-HCR-target_3 | tctcccttgaaactctataatt***AAA***CGTCGGAGTGGGTTCAGTCTA | 360 | 32.00% |
| BTV1-WT-S1_RNA-HCR-target_4 | ttaatgggtatatctccgtata***AAA***CGTCGGAGTGGGTTCAGTCTA | 433 | 32.00% |
| BTV1-WT-S1_RNA-HCR-target_5 | gcactcagttcattgatgaaac***AAA***CGTCGGAGTGGGTTCAGTCTA | 466 | 41.00% |
| BTV1-WT-S1_RNA-HCR-target_6 | tggaatggtgaatcgaaacggt***AAA***CGTCGGAGTGGGTTCAGTCTA | 538 | 45.00% |
| BTV1-WT-S1_RNA-HCR-target_7 | ttacaaaccaagagtatggtgc***AAA***CGTCGGAGTGGGTTCAGTCTA | 798 | 41.00% |
| BTV1-WT-S1_RNA-HCR-target_8 | aaacacatcctttgaattcctg***AAA***CGTCGGAGTGGGTTCAGTCTA | 878 | 36.00% |
| BTV1-WT-S1_RNA-HCR-target_9 | aatagggctttgtatcgatttg***AAA***CGTCGGAGTGGGTTCAGTCTA | 922 | 36.00% |
| BTV1-WT-S1_RNA-HCR-target_10 | aatggtgtagtataaaccgtct***AAA***CGTCGGAGTGGGTTCAGTCTA | 1066 | 36.00% |
| BTV1-WT-S1_RNA-HCR-target_11 | cttggaccaaatctctttttta***AAA***CGTCGGAGTGGGTTCAGTCTA | 1384 | 32.00% |
| BTV1-WT-S1_RNA-HCR-target_12 | aagccttaatccttgaattgat***AAA***CGTCGGAGTGGGTTCAGTCTA | 1431 | 32.00% |
| BTV1-WT-S1_RNA-HCR-target_13 | tcagtgaaaacagtatgtcctt***AAA***CGTCGGAGTGGGTTCAGTCTA | 1468 | 36.00% |
| BTV1-WT-S1_RNA-HCR-target_14 | agtttggtatagttctacacta***AAA***CGTCGGAGTGGGTTCAGTCTA | 1514 | 32.00% |
| BTV1-WT-S1_RNA-HCR-target_15 | caattgtgaatatatcgcggtc***AAA***CGTCGGAGTGGGTTCAGTCTA | 1767 | 41.00% |
| BTV1-WT-S1_RNA-HCR-target_16 | gcataccggttcgaaaattatg***AAA***CGTCGGAGTGGGTTCAGTCTA | 1827 | 41.00% |
| BTV1-WT-S1_RNA-HCR-target_17 | tgtagttttaaacagtcttcgc***AAA***CGTCGGAGTGGGTTCAGTCTA | 1973 | 36.00% |
| BTV1-WT-S1_RNA-HCR-target_18 | caatactcgatactggaagcac***AAA***CGTCGGAGTGGGTTCAGTCTA | 2061 | 45.00% |
| BTV1-WT-S1_RNA-HCR-target_19 | attaaaaattgaagccgccact***AAA***CGTCGGAGTGGGTTCAGTCTA | 2357 | 36.00% |
| BTV1-WT-S1_RNA-HCR-target_20 | taagcattaatattagctgcgc***AAA***CGTCGGAGTGGGTTCAGTCTA | 2607 | 36.00% |
| BTV1-WT-S1_RNA-HCR-target_21 | gaatctgtatcaacgtaaaccc***AAA***CGTCGGAGTGGGTTCAGTCTA | 2721 | 41.00% |
| BTV1-WT-S1_RNA-HCR-target_22 | cataacgatattaagagctgcc***AAA***CGTCGGAGTGGGTTCAGTCTA | 2798 | 41.00% |
| BTV1-WT-S1_RNA-HCR-target_23 | tgatattctagttcgtaaccac***AAA***CGTCGGAGTGGGTTCAGTCTA | 3136 | 36.00% |
| BTV1-WT-S1_RNA-HCR-target_24 | agatgttgatattactcctgac***AAA***CGTCGGAGTGGGTTCAGTCTA | 3209 | 36.00% |
| BTV1-WT-S1_RNA-HCR-target_25 | ccttcgaagtatacgtcaaaga***AAA***CGTCGGAGTGGGTTCAGTCTA | 3241 | 41.00% |
| BTV1-WT-S1_RNA-HCR-target_26 | gataagttctggtctggaaaca***AAA***CGTCGGAGTGGGTTCAGTCTA | 3286 | 41.00% |
| BTV1-WT-S1_RNA-HCR-target_27 | caatatcacatcaatccgatct***AAA***CGTCGGAGTGGGTTCAGTCTA | 3377 | 36.00% |
| BTV1-WT-S1_RNA-HCR-target_28 | tctcaattacgttcagaatggt***AAA***CGTCGGAGTGGGTTCAGTCTA | 3438 | 36.00% |
| BTV1-WT-S1_RNA-HCR-target_29 | ttttcgtcaactgatagggata***AAA***CGTCGGAGTGGGTTCAGTCTA | 3672 | 36.00% |
| BTV1-WT-S1_RNA-HCR-target_30 | actttaattttaggtacgtgcg***AAA***CGTCGGAGTGGGTTCAGTCTA | 3835 | 36.00% |

| S6-HCR-probes-cy3 | Probe (5'-> 3') | Probe position | Percent GC |
| --- | --- | --- | --- |
| RNA-HCR-Probe-H1_5'-cy3 | cy3 - 5' - AGTACATGTCGTGGTGGTAGC***TTGTATGAA***GCTACCACCACG - 3' |  |  |
| RNA-HCR-Probe-H2_3'-cy3 | 5' - GCTACCACCACG***ACATGTACT***CGTGGTGGTAGCTTCATACAA - 3' - cy3 |  |  |
| BTV1-WT-S6_RNA-HCR-target_1 | gtggttgccaactagagaac***AAA***GCTACCACCACGACATGTACT | 10 | 50.00% |
| BTV1-WT-S6_RNA-HCR-target_2 | gccaaaaaagttctcgtggc***AAA***GCTACCACCACGACATGTACT | 83 | 50.00% |
| BTV1-WT-S6_RNA-HCR-target_3 | atgactgcaagtccattgtg***AAA***GCTACCACCACGACATGTACT | 111 | 45.00% |
| BTV1-WT-S6_RNA-HCR-target_4 | aatttatatgctttcgccgg***AAA***GCTACCACCACGACATGTACT | 215 | 40.00% |
| BTV1-WT-S6_RNA-HCR-target_5 | ttttgaagcattggagccag***AAA***GCTACCACCACGACATGTACT | 277 | 45.00% |
| BTV1-WT-S6_RNA-HCR-target_6 | cggtaatcttcgagcagttg***AAA***GCTACCACCACGACATGTACT | 353 | 50.00% |
| BTV1-WT-S6_RNA-HCR-target_7 | taaccaatgcggattgcttc***AAA***GCTACCACCACGACATGTACT | 400 | 45.00% |
| BTV1-WT-S6_RNA-HCR-target_8 | gttggattcacgatttgacc***AAA***GCTACCACCACGACATGTACT | 488 | 45.00% |
| BTV1-WT-S6_RNA-HCR-target_9 | gcatccgggttgtagaaata***AAA***GCTACCACCACGACATGTACT | 539 | 45.00% |
| BTV1-WT-S6_RNA-HCR-target_10 | tctcaacctcacgtttgatt***AAA***GCTACCACCACGACATGTACT | 613 | 40.00% |
| BTV1-WT-S6_RNA-HCR-target_11 | cagtgtagggacatgtgtta***AAA***GCTACCACCACGACATGTACT | 640 | 45.00% |
| BTV1-WT-S6_RNA-HCR-target_12 | gatcagctgaatcggcaaga***AAA***GCTACCACCACGACATGTACT | 690 | 50.00% |
| BTV1-WT-S6_RNA-HCR-target_13 | aatattgctgtatcgccatc***AAA***GCTACCACCACGACATGTACT | 760 | 40.00% |
| BTV1-WT-S6_RNA-HCR-target_14 | atctgacctcttcagcataa***AAA***GCTACCACCACGACATGTACT | 790 | 40.00% |
| BTV1-WT-S6_RNA-HCR-target_15 | cgttggaacccttccaaaaa***AAA***GCTACCACCACGACATGTACT | 822 | 45.00% |
| BTV1-WT-S6_RNA-HCR-target_16 | gttgggaaatcgcgtcttat***AAA***GCTACCACCACGACATGTACT | 872 | 45.00% |
| BTV1-WT-S6_RNA-HCR-target_17 | taatcataggtagcagccag***AAA***GCTACCACCACGACATGTACT | 952 | 45.00% |
| BTV1-WT-S6_RNA-HCR-target_18 | gctgacatgtatgctttcta***AAA***GCTACCACCACGACATGTACT | 1030 | 40.00% |
| BTV1-WT-S6_RNA-HCR-target_19 | gatctgctttgagtgtttca***AAA***GCTACCACCACGACATGTACT | 1059 | 40.00% |
| BTV1-WT-S6_RNA-HCR-target_20 | tgatgcgcgtacatcaatca***AAA***GCTACCACCACGACATGTACT | 1092 | 45.00% |
| BTV1-WT-S6_RNA-HCR-target_21 | gaatcatttcccacatgttc***AAA***GCTACCACCACGACATGTACT | 1154 | 40.00% |
| BTV1-WT-S6_RNA-HCR-target_22 | cggtgaatgtgaatcgcagt***AAA***GCTACCACCACGACATGTACT | 1274 | 50.00% |
| BTV1-WT-S6_RNA-HCR-target_23 | cgcgtccgagcatgaaaata***AAA***GCTACCACCACGACATGTACT | 1342 | 50.00% |
| BTV1-WT-S6_RNA-HCR-target_24 | ccgtagcacacaaagcagaa***AAA***GCTACCACCACGACATGTACT | 1415 | 50.00% |
| BTV1-WT-S6_RNA-HCR-target_25 | tccggtcctttcaaaatgat***AAA***GCTACCACCACGACATGTACT | 1496 | 40.00% |
| BTV1-WT-S6_RNA-HCR-target_26 | aaacatcgtagcataagccc***AAA***GCTACCACCACGACATGTACT | 1539 | 45.00% |
| BTV1-WT-S6_RNA-HCR-target_27 | gggtgataatgcatcgaacc***AAA***GCTACCACCACGACATGTACT | 1564 | 50.00% |
| BTV1-WT-S6_RNA-HCR-target_28 | cagcgaagtgaaccttttct***AAA***GCTACCACCACGACATGTACT | 1594 | 45.00% |
| BTV1-WT-S6_RNA-HCR-target_29 | agcgagattgataacttccc***AAA***GCTACCACCACGACATGTACT | 1641 | 45.00% |
| BTV1-WT-S6_RNA-HCR-target_30 | aatactccatccacatctga***AAA***GCTACCACCACGACATGTACT | 1672 | 40.00% |

| S10-HCR-probes-cy5 | Probe (5'-> 3') | Probe position | Percent GC |
| --- | --- | --- | --- |
| RNA-HCR-Probe-H1_5'-cy5 | cy5 - 5' - TGTTGCAAAGGAACGTCGAGC***TGTAATGGT***GCTCGACGTTCC - 3' |  |  |
| RNA-HCR-Probe-H2_3'-cy5 | 5' - GCTCGACGTTCC***TTTGCAACA***GGAACGTCGAGCACCATTACA - 3' - cy5 |  |  |
| BTV1-WT-S10_RNA-HCR-target_1 | atggcagcgacacttttt***AAA***GCTCGACGTTCCTTTGCAACA | 4 | 44.00% |
| BTV1-WT-S10_RNA-HCR-target_2 | tttggatcagcccggata***AAA***GCTCGACGTTCCTTTGCAACA | 24 | 50.00% |
| BTV1-WT-S10_RNA-HCR-target_3 | tcattttttcttcctcga***AAA***GCTCGACGTTCCTTTGCAACA | 45 | 33.00% |
| BTV1-WT-S10_RNA-HCR-target_4 | ttcaacccgttcctgatt***AAA***GCTCGACGTTCCTTTGCAACA | 68 | 44.00% |
| BTV1-WT-S10_RNA-HCR-target_5 | acacggaccagactcagc***AAA***GCTCGACGTTCCTTTGCAACA | 88 | 61.00% |
| BTV1-WT-S10_RNA-HCR-target_6 | gcggttgggaaatcgtgt***AAA***GCTCGACGTTCCTTTGCAACA | 111 | 56.00% |
| BTV1-WT-S10_RNA-HCR-target_7 | cgcactcggagcatatct***AAA***GCTCGACGTTCCTTTGCAACA | 131 | 56.00% |
| BTV1-WT-S10_RNA-HCR-target_8 | aggcatcgacgatggcat***AAA***GCTCGACGTTCCTTTGCAACA | 152 | 56.00% |
| BTV1-WT-S10_RNA-HCR-target_9 | gctttgtccaagatttca***AAA***GCTCGACGTTCCTTTGCAACA | 181 | 39.00% |
| BTV1-WT-S10_RNA-HCR-target_10 | cacccgttgtatttgaca***AAA***GCTCGACGTTCCTTTGCAACA | 201 | 44.00% |
| BTV1-WT-S10_RNA-HCR-target_11 | cgctttttgtgtttgcgt***AAA***GCTCGACGTTCCTTTGCAACA | 221 | 44.00% |
| BTV1-WT-S10_RNA-HCR-target_12 | gatgcgaatgcagccttc***AAA***GCTCGACGTTCCTTTGCAACA | 241 | 56.00% |
| BTV1-WT-S10_RNA-HCR-target_13 | cacgaaacgcttctgcgt***AAA***GCTCGACGTTCCTTTGCAACA | 261 | 56.00% |
| BTV1-WT-S10_RNA-HCR-target_14 | ctgtctcaacctcacgtc***AAA***GCTCGACGTTCCTTTGCAACA | 281 | 56.00% |
| BTV1-WT-S10_RNA-HCR-target_15 | ctttaagcctcctaggtc***AAA***GCTCGACGTTCCTTTGCAACA | 344 | 50.00% |
| BTV1-WT-S10_RNA-HCR-target_16 | aacaaccgccgctatcag***AAA***GCTCGACGTTCCTTTGCAACA | 392 | 56.00% |
| BTV1-WT-S10_RNA-HCR-target_17 | agcttaaacgccacgctc***AAA***GCTCGACGTTCCTTTGCAACA | 448 | 56.00% |
| BTV1-WT-S10_RNA-HCR-target_18 | ccattgaggtatctcagc***AAA***GCTCGACGTTCCTTTGCAACA | 479 | 50.00% |
| BTV1-WT-S10_RNA-HCR-target_19 | cattgggttcagactctt***AAA***GCTCGACGTTCCTTTGCAACA | 500 | 44.00% |
| BTV1-WT-S10_RNA-HCR-target_20 | cccaaattcaccacacct***AAA***GCTCGACGTTCCTTTGCAACA | 520 | 50.00% |
| BTV1-WT-S10_RNA-HCR-target_21 | gaccatcatcaggaaggt***AAA***GCTCGACGTTCCTTTGCAACA | 542 | 50.00% |
| BTV1-WT-S10_RNA-HCR-target_22 | cttctctcgctttttgcg***AAA***GCTCGACGTTCCTTTGCAACA | 562 | 50.00% |
| BTV1-WT-S10_RNA-HCR-target_23 | tgtcaatttgctggttca***AAA***GCTCGACGTTCCTTTGCAACA | 582 | 39.00% |
| BTV1-WT-S10_RNA-HCR-target_24 | tcttcatcacttccttct***AAA***GCTCGACGTTCCTTTGCAACA | 606 | 39.00% |
| BTV1-WT-S10_RNA-HCR-target_25 | tcctcaccgcatcattat***AAA***GCTCGACGTTCCTTTGCAACA | 633 | 44.00% |
| BTV1-WT-S10_RNA-HCR-target_26 | ccatctagcgggactgat***AAA***GCTCGACGTTCCTTTGCAACA | 670 | 56.00% |
| BTV1-WT-S10_RNA-HCR-target_27 | gccactctacctactgat***AAA***GCTCGACGTTCCTTTGCAACA | 711 | 50.00% |
| BTV1-WT-S10_RNA-HCR-target_28 | acacgatgcagacctcgg***AAA***GCTCGACGTTCCTTTGCAACA | 732 | 61.00% |
| BTV1-WT-S10_RNA-HCR-target_29 | gtctgcatcgtgagatca***AAA***GCTCGACGTTCCTTTGCAACA | 758 | 50.00% |
| BTV1-WT-S10_RNA-HCR-target_30 | cccgttagacagcagtag***AAA***GCTCGACGTTCCTTTGCAACA | 778 | 56.00% |

| S6-HCR-probes-cy5 | Probe (5'-> 3') | Probe position | Percent GC |
| --- | --- | --- | --- |
| RNA-HCR-Probe-H1_5'-cy5 | cy5 - 5' - TGTTGCAAAGGAACGTCGAGC***TGTAATGGT***GCTCGACGTTCC - 3' |  |  |
| RNA-HCR-Probe-H2_3'-cy5 | 5' - GCTCGACGTTCC***TTTGCAACA***GGAACGTCGAGCACCATTACA - 3' - cy5 |  |  |
| BTV1-WT-S6_RNA-HCR-target_1 | gtggttgccaactagagaac***AAA***GCTCGACGTTCCTTTGCAACA | 10 | 50.00% |
| BTV1-WT-S6_RNA-HCR-target_2 | gccaaaaaagttctcgtggc***AAA***GCTCGACGTTCCTTTGCAACA | 83 | 50.00% |
| BTV1-WT-S6_RNA-HCR-target_3 | atgactgcaagtccattgtg***AAA***GCTCGACGTTCCTTTGCAACA | 111 | 45.00% |
| BTV1-WT-S6_RNA-HCR-target_4 | aatttatatgctttcgccgg***AAA***GCTCGACGTTCCTTTGCAACA | 215 | 40.00% |
| BTV1-WT-S6_RNA-HCR-target_5 | ttttgaagcattggagccag***AAA***GCTCGACGTTCCTTTGCAACA | 277 | 45.00% |
| BTV1-WT-S6_RNA-HCR-target_6 | cggtaatcttcgagcagttg***AAA***GCTCGACGTTCCTTTGCAACA | 353 | 50.00% |
| BTV1-WT-S6_RNA-HCR-target_7 | taaccaatgcggattgcttc***AAA***GCTCGACGTTCCTTTGCAACA | 400 | 45.00% |
| BTV1-WT-S6_RNA-HCR-target_8 | gttggattcacgatttgacc***AAA***GCTCGACGTTCCTTTGCAACA | 488 | 45.00% |
| BTV1-WT-S6_RNA-HCR-target_9 | gcatccgggttgtagaaata***AAA***GCTCGACGTTCCTTTGCAACA | 539 | 45.00% |
| BTV1-WT-S6_RNA-HCR-target_10 | tctcaacctcacgtttgatt***AAA***GCTCGACGTTCCTTTGCAACA | 613 | 40.00% |
| BTV1-WT-S6_RNA-HCR-target_11 | cagtgtagggacatgtgtta***AAA***GCTCGACGTTCCTTTGCAACA | 640 | 45.00% |
| BTV1-WT-S6_RNA-HCR-target_12 | gatcagctgaatcggcaaga***AAA***GCTCGACGTTCCTTTGCAACA | 690 | 50.00% |
| BTV1-WT-S6_RNA-HCR-target_13 | aatattgctgtatcgccatc***AAA***GCTCGACGTTCCTTTGCAACA | 760 | 40.00% |
| BTV1-WT-S6_RNA-HCR-target_14 | atctgacctcttcagcataa***AAA***GCTCGACGTTCCTTTGCAACA | 790 | 40.00% |
| BTV1-WT-S6_RNA-HCR-target_15 | cgttggaacccttccaaaaa***AAA***GCTCGACGTTCCTTTGCAACA | 822 | 45.00% |
| BTV1-WT-S6_RNA-HCR-target_16 | gttgggaaatcgcgtcttat***AAA***GCTCGACGTTCCTTTGCAACA | 872 | 45.00% |
| BTV1-WT-S6_RNA-HCR-target_17 | taatcataggtagcagccag***AAA***GCTCGACGTTCCTTTGCAACA | 952 | 45.00% |
| BTV1-WT-S6_RNA-HCR-target_18 | gctgacatgtatgctttcta***AAA***GCTCGACGTTCCTTTGCAACA | 1030 | 40.00% |
| BTV1-WT-S6_RNA-HCR-target_19 | gatctgctttgagtgtttca***AAA***GCTCGACGTTCCTTTGCAACA | 1059 | 40.00% |
| BTV1-WT-S6_RNA-HCR-target_20 | tgatgcgcgtacatcaatca***AAA***GCTCGACGTTCCTTTGCAACA | 1092 | 45.00% |
| BTV1-WT-S6_RNA-HCR-target_21 | gaatcatttcccacatgttc***AAA***GCTCGACGTTCCTTTGCAACA | 1154 | 40.00% |
| BTV1-WT-S6_RNA-HCR-target_22 | cggtgaatgtgaatcgcagt***AAA***GCTCGACGTTCCTTTGCAACA | 1274 | 50.00% |
| BTV1-WT-S6_RNA-HCR-target_23 | cgcgtccgagcatgaaaata***AAA***GCTCGACGTTCCTTTGCAACA | 1342 | 50.00% |
| BTV1-WT-S6_RNA-HCR-target_24 | ccgtagcacacaaagcagaa***AAA***GCTCGACGTTCCTTTGCAACA | 1415 | 50.00% |
| BTV1-WT-S6_RNA-HCR-target_25 | tccggtcctttcaaaatgat***AAA***GCTCGACGTTCCTTTGCAACA | 1496 | 40.00% |
| BTV1-WT-S6_RNA-HCR-target_26 | aaacatcgtagcataagccc***AAA***GCTCGACGTTCCTTTGCAACA | 1539 | 45.00% |
| BTV1-WT-S6_RNA-HCR-target_27 | gggtgataatgcatcgaacc***AAA***GCTCGACGTTCCTTTGCAACA | 1564 | 50.00% |
| BTV1-WT-S6_RNA-HCR-target_28 | cagcgaagtgaaccttttct***AAA***GCTCGACGTTCCTTTGCAACA | 1594 | 45.00% |
| BTV1-WT-S6_RNA-HCR-target_29 | agcgagattgataacttccc***AAA***GCTCGACGTTCCTTTGCAACA | 1641 | 45.00% |
| BTV1-WT-S6_RNA-HCR-target_30 | aatactccatccacatctga***AAA***GCTCGACGTTCCTTTGCAACA | 1672 | 40.00% |

| S1-HCR-probes-cy5 | Probe (5'-> 3') | Probe position | Percent GC |
| --- | --- | --- | --- |
| RNA-HCR-Probe-H1_5'-cy5 | cy5 - 5' - TGTTGCAAAGGAACGTCGAGC***TGTAATGGT***GCTCGACGTTCC - 3' |  |  |
| RNA-HCR-Probe-H2_3'-cy5 | 5' - GCTCGACGTTCC***TTTGCAACA***GGAACGTCGAGCACCATTACA - 3' - cy5 |  |  |
| BTV1-WT-S1_RNA-HCR-target_1 | aacctcatctaatactggtatc***AAA***GCTCGACGTTCCTTTGCAACA | 227 | 36.00% |
| BTV1-WT-S1_RNA-HCR-target_2 | cctcaaaagcgtaactttggaa***AAA***GCTCGACGTTCCTTTGCAACA | 279 | 41.00% |
| BTV1-WT-S1_RNA-HCR-target_3 | tctcccttgaaactctataatt***AAA***GCTCGACGTTCCTTTGCAACA | 360 | 32.00% |
| BTV1-WT-S1_RNA-HCR-target_4 | ttaatgggtatatctccgtata***AAA***GCTCGACGTTCCTTTGCAACA | 433 | 32.00% |
| BTV1-WT-S1_RNA-HCR-target_5 | gcactcagttcattgatgaaac***AAA***GCTCGACGTTCCTTTGCAACA | 466 | 41.00% |
| BTV1-WT-S1_RNA-HCR-target_6 | tggaatggtgaatcgaaacggt***AAA***GCTCGACGTTCCTTTGCAACA | 538 | 45.00% |
| BTV1-WT-S1_RNA-HCR-target_7 | ttacaaaccaagagtatggtgc***AAA***GCTCGACGTTCCTTTGCAACA | 798 | 41.00% |
| BTV1-WT-S1_RNA-HCR-target_8 | aaacacatcctttgaattcctg***AAA***GCTCGACGTTCCTTTGCAACA | 878 | 36.00% |
| BTV1-WT-S1_RNA-HCR-target_9 | aatagggctttgtatcgatttg***AAA***GCTCGACGTTCCTTTGCAACA | 922 | 36.00% |
| BTV1-WT-S1_RNA-HCR-target_10 | aatggtgtagtataaaccgtct***AAA***GCTCGACGTTCCTTTGCAACA | 1066 | 36.00% |
| BTV1-WT-S1_RNA-HCR-target_11 | cttggaccaaatctctttttta***AAA***GCTCGACGTTCCTTTGCAACA | 1384 | 32.00% |
| BTV1-WT-S1_RNA-HCR-target_12 | aagccttaatccttgaattgat***AAA***GCTCGACGTTCCTTTGCAACA | 1431 | 32.00% |
| BTV1-WT-S1_RNA-HCR-target_13 | tcagtgaaaacagtatgtcctt***AAA***GCTCGACGTTCCTTTGCAACA | 1468 | 36.00% |
| BTV1-WT-S1_RNA-HCR-target_14 | agtttggtatagttctacacta***AAA***GCTCGACGTTCCTTTGCAACA | 1514 | 32.00% |
| BTV1-WT-S1_RNA-HCR-target_15 | caattgtgaatatatcgcggtc***AAA***GCTCGACGTTCCTTTGCAACA | 1767 | 41.00% |
| BTV1-WT-S1_RNA-HCR-target_16 | gcataccggttcgaaaattatg***AAA***GCTCGACGTTCCTTTGCAACA | 1827 | 41.00% |
| BTV1-WT-S1_RNA-HCR-target_17 | tgtagttttaaacagtcttcgc***AAA***GCTCGACGTTCCTTTGCAACA | 1973 | 36.00% |
| BTV1-WT-S1_RNA-HCR-target_18 | caatactcgatactggaagcac***AAA***GCTCGACGTTCCTTTGCAACA | 2061 | 45.00% |
| BTV1-WT-S1_RNA-HCR-target_19 | attaaaaattgaagccgccact***AAA***GCTCGACGTTCCTTTGCAACA | 2357 | 36.00% |
| BTV1-WT-S1_RNA-HCR-target_20 | taagcattaatattagctgcgc***AAA***GCTCGACGTTCCTTTGCAACA | 2607 | 36.00% |
| BTV1-WT-S1_RNA-HCR-target_21 | gaatctgtatcaacgtaaaccc***AAA***GCTCGACGTTCCTTTGCAACA | 2721 | 41.00% |
| BTV1-WT-S1_RNA-HCR-target_22 | cataacgatattaagagctgcc***AAA***GCTCGACGTTCCTTTGCAACA | 2798 | 41.00% |
| BTV1-WT-S1_RNA-HCR-target_23 | tgatattctagttcgtaaccac***AAA***GCTCGACGTTCCTTTGCAACA | 3136 | 36.00% |
| BTV1-WT-S1_RNA-HCR-target_24 | agatgttgatattactcctgac***AAA***GCTCGACGTTCCTTTGCAACA | 3209 | 36.00% |
| BTV1-WT-S1_RNA-HCR-target_25 | ccttcgaagtatacgtcaaaga***AAA***GCTCGACGTTCCTTTGCAACA | 3241 | 41.00% |
| BTV1-WT-S1_RNA-HCR-target_26 | gataagttctggtctggaaaca***AAA***GCTCGACGTTCCTTTGCAACA | 3286 | 41.00% |
| BTV1-WT-S1_RNA-HCR-target_27 | caatatcacatcaatccgatct***AAA***GCTCGACGTTCCTTTGCAACA | 3377 | 36.00% |
| BTV1-WT-S1_RNA-HCR-target_28 | tctcaattacgttcagaatggt***AAA***GCTCGACGTTCCTTTGCAACA | 3438 | 36.00% |
| BTV1-WT-S1_RNA-HCR-target_29 | ttttcgtcaactgatagggata***AAA***GCTCGACGTTCCTTTGCAACA | 3672 | 36.00% |
| BTV1-WT-S1_RNA-HCR-target_30 | actttaattttaggtacgtgcg***AAA***GCTCGACGTTCCTTTGCAACA | 3835 | 36.00% |

| ActB-HCR-probes-Cy5 | Probe (5'-> 3') | Probe position | Percent GC |
| --- | --- | --- | --- |
| RNA-HCR-Probe-H1_5'-cy5 | cy5 - 5' - TGTTGCAAAGGAACGTCGAGC***TGTAATGGT***GCTCGACGTTCC - 3' |  |  |
| RNA-HCR-Probe-H2_3'-cy5 | 5' - GCTCGACGTTCC***TTTGCAACA***GGAACGTCGAGCACCATTACA - 3' - cy5 |  |  |
| BHK-21_ actb-mRNA-HCR-target_1 | acaacgagcgcagcgatatc**AAA**GCTCGACGTTCCTTTGCAACA | 10 | 55.00% |
| BHK-21_ actb-mRNA-HCR-target_2 | cacgatggaggggaagacgg**AAA**GCTCGACGTTCCTTTGCAACA | 86 | 65.00% |
| BHK-21_ actb-mRNA-HCR-target_3 | agaatacctctcttgctctg**AAA**GCTCGACGTTCCTTTGCAACA | 175 | 45.00% |
| BHK-21_ actb-mRNA-HCR-target_4 | gtgacaatgccgtgttcaat**AAA**GCTCGACGTTCCTTTGCAACA | 211 | 45.00% |
| BHK-21_ actb-mRNA-HCR-target_5 | cagatcttctccatatcgtc**AAA**GCTCGACGTTCCTTTGCAACA | 238 | 45.00% |
| BHK-21_ actb-mRNA-HCR-target_6 | cacgcagctcgttgtagaag**AAA**GCTCGACGTTCCTTTGCAACA | 267 | 55.00% |
| BHK-21_ actb-mRNA-HCR-target_7 | tgggtcatcttttcacggtt**AAA**GCTCGACGTTCCTTTGCAACA | 343 | 45.00% |
| BHK-21_ actb-mRNA-HCR-target_8 | ggtgttgaaggtctcaaaca**AAA**GCTCGACGTTCCTTTGCAACA | 368 | 45.00% |
| BHK-21_ actb-mRNA-HCR-target_9 | cctgaatggctacgtacatg**AAA**GCTCGACGTTCCTTTGCAACA | 393 | 50.00% |
| BHK-21_ actb-mRNA-HCR-target_10 | gtacgaccagaggcatacag**AAA**GCTCGACGTTCCTTTGCAACA | 424 | 55.00% |
| BHK-21_ actb-mRNA-HCR-target_11 | ctccggagtccatcacaatg**AAA**GCTCGACGTTCCTTTGCAACA | 450 | 55.00% |
| BHK-21_ actb-mRNA-HCR-target_12 | tcatagatgggcacagtgtg**AAA**GCTCGACGTTCCTTTGCAACA | 481 | 50.00% |
| BHK-21_ actb-mRNA-HCR-target_13 | caggatggcatgagggagag**AAA**GCTCGACGTTCCTTTGCAACA | 509 | 60.00% |
| BHK-21_ actb-mRNA-HCR-target_14 | atcttcatgaggtagtctgt**AAA**GCTCGACGTTCCTTTGCAACA | 556 | 40.00% |
| BHK-21_ actb-mRNA-HCR-target_15 | ctgtggtggtgaagctgtag**AAA**GCTCGACGTTCCTTTGCAACA | 591 | 55.00% |
| BHK-21_ actb-mRNA-HCR-target_16 | ctctttgatgtcacgcacaa**AAA**GCTCGACGTTCCTTTGCAACA | 623 | 45.00% |
| BHK-21_ actb-mRNA-HCR-target_17 | cgaagtccagggcaacatag**AAA**GCTCGACGTTCCTTTGCAACA | 651 | 55.00% |
| BHK-21_ actb-mRNA-HCR-target_18 | ctccagggaggaagaggatg**AAA**GCTCGACGTTCCTTTGCAACA | 692 | 60.00% |
| BHK-21_ actb-mRNA-HCR-target_19 | tcgttgccaatggtgatgac**AAA**GCTCGACGTTCCTTTGCAACA | 739 | 50.00% |
| BHK-21_ actb-mRNA-HCR-target_20 | gaaaagggcctcagggcaac**AAA**GCTCGACGTTCCTTTGCAACA | 767 | 60.00% |
| BHK-21_ actb-mRNA-HCR-target_21 | attccatacccaggaaggaa**AAA**GCTCGACGTTCCTTTGCAACA | 792 | 45.00% |
| BHK-21_ actb-mRNA-HCR-target_22 | aatgtagtttcgtggatgcc**AAA**GCTCGACGTTCCTTTGCAACA | 817 | 45.00% |
| BHK-21_ actb-mRNA-HCR-target_23 | gacgtcacacttcatgatgg**AAA**GCTCGACGTTCCTTTGCAACA | 842 | 50.00% |
| BHK-21_ actb-mRNA-HCR-target_24 | tgttggcatagaggtctttg**AAA**GCTCGACGTTCCTTTGCAACA | 870 | 45.00% |
| BHK-21_ actb-mRNA-HCR-target_25 | caatgcctgggtacatggtg**AAA**GCTCGACGTTCCTTTGCAACA | 909 | 55.00% |
| BHK-21_ actb-mRNA-HCR-target_26 | agagcagtgatctccttctg**AAA**GCTCGACGTTCCTTTGCAACA | 940 | 50.00% |
| BHK-21_ actb-mRNA-HCR-target_27 | ggggagcaatgatcttgatc**AAA**GCTCGACGTTCCTTTGCAACA | 978 | 50.00% |
| BHK-21_ actb-mRNA-HCR-target_28 | cgatccacacagagtacttg**AAA**GCTCGACGTTCCTTTGCAACA | 1005 | 50.00% |
| BHK-21_ actb-mRNA-HCR-target_29 | tgatccacatctgctggaag**AAA**GCTCGACGTTCCTTTGCAACA | 1053 | 50.00% |
| BHK-21_ actb-mRNA-HCR-target_30 | tagaagcatttgcggtggac**AAA**GCTCGACGTTCCTTTGCAACA | 1108 | 50.00% |

| ActB-HCR-probes-Cy3 | Probe (5'-> 3') | Probe position | Percent GC |
| --- | --- | --- | --- |
| RNA-HCR-Probe-H1_5'-cy3 | cy3 - 5' - AGTACATGTCGTGGTGGTAGC***TTGTATGAA***GCTACCACCACG - 3' |  |  |
| RNA-HCR-Probe-H2_3'-cy3 | 5' - GCTACCACCACG***ACATGTACT***CGTGGTGGTAGCTTCATACAA - 3' - cy3 |  |  |
| BHK-21_ actb-mRNA-HCR-target_1 | acaacgagcgcagcgatatc**AAA**GCTACCACCACGACATGTACT | 10 | 55.00% |
| BHK-21_ actb-mRNA-HCR-target_2 | cacgatggaggggaagacgg**AAA**GCTACCACCACGACATGTACT | 86 | 65.00% |
| BHK-21_ actb-mRNA-HCR-target_3 | agaatacctctcttgctctg**AAA**GCTACCACCACGACATGTACT | 175 | 45.00% |
| BHK-21_ actb-mRNA-HCR-target_4 | gtgacaatgccgtgttcaat**AAA**GCTACCACCACGACATGTACT | 211 | 45.00% |
| BHK-21_ actb-mRNA-HCR-target_5 | cagatcttctccatatcgtc**AAA**GCTACCACCACGACATGTACT | 238 | 45.00% |
| BHK-21_ actb-mRNA-HCR-target_6 | cacgcagctcgttgtagaag**AAA**GCTACCACCACGACATGTACT | 267 | 55.00% |
| BHK-21_ actb-mRNA-HCR-target_7 | tgggtcatcttttcacggtt**AAA**GCTACCACCACGACATGTACT | 343 | 45.00% |
| BHK-21_ actb-mRNA-HCR-target_8 | ggtgttgaaggtctcaaaca**AAA**GCTACCACCACGACATGTACT | 368 | 45.00% |
| BHK-21_ actb-mRNA-HCR-target_9 | cctgaatggctacgtacatg**AAA**GCTACCACCACGACATGTACT | 393 | 50.00% |
| BHK-21_ actb-mRNA-HCR-target_10 | gtacgaccagaggcatacag**AAA**GCTACCACCACGACATGTACT | 424 | 55.00% |
| BHK-21_ actb-mRNA-HCR-target_11 | ctccggagtccatcacaatg**AAA**GCTACCACCACGACATGTACT | 450 | 55.00% |
| BHK-21_ actb-mRNA-HCR-target_12 | tcatagatgggcacagtgtg**AAA**GCTACCACCACGACATGTACT | 481 | 50.00% |
| BHK-21_ actb-mRNA-HCR-target_13 | caggatggcatgagggagag**AAA**GCTACCACCACGACATGTACT | 509 | 60.00% |
| BHK-21_ actb-mRNA-HCR-target_14 | atcttcatgaggtagtctgt**AAA**GCTACCACCACGACATGTACT | 556 | 40.00% |
| BHK-21_ actb-mRNA-HCR-target_15 | ctgtggtggtgaagctgtag**AAA**GCTACCACCACGACATGTACT | 591 | 55.00% |
| BHK-21_ actb-mRNA-HCR-target_16 | ctctttgatgtcacgcacaa**AAA**GCTACCACCACGACATGTACT | 623 | 45.00% |
| BHK-21_ actb-mRNA-HCR-target_17 | cgaagtccagggcaacatag**AAA**GCTACCACCACGACATGTACT | 651 | 55.00% |
| BHK-21_ actb-mRNA-HCR-target_18 | ctccagggaggaagaggatg**AAA**GCTACCACCACGACATGTACT | 692 | 60.00% |
| BHK-21_ actb-mRNA-HCR-target_19 | tcgttgccaatggtgatgac**AAA**GCTACCACCACGACATGTACT | 739 | 50.00% |
| BHK-21_ actb-mRNA-HCR-target_20 | gaaaagggcctcagggcaac**AAA**GCTACCACCACGACATGTACT | 767 | 60.00% |
| BHK-21_ actb-mRNA-HCR-target_21 | attccatacccaggaaggaa**AAA**GCTACCACCACGACATGTACT | 792 | 45.00% |
| BHK-21_ actb-mRNA-HCR-target_22 | aatgtagtttcgtggatgcc**AAA**GCTACCACCACGACATGTACT | 817 | 45.00% |
| BHK-21_ actb-mRNA-HCR-target_23 | gacgtcacacttcatgatgg**AAA**GCTACCACCACGACATGTACT | 842 | 50.00% |
| BHK-21_ actb-mRNA-HCR-target_24 | tgttggcatagaggtctttg**AAA**GCTACCACCACGACATGTACT | 870 | 45.00% |
| BHK-21_ actb-mRNA-HCR-target_25 | caatgcctgggtacatggtg**AAA**GCTACCACCACGACATGTACT | 909 | 55.00% |
| BHK-21_ actb-mRNA-HCR-target_26 | agagcagtgatctccttctg**AAA**GCTACCACCACGACATGTACT | 940 | 50.00% |
| BHK-21_ actb-mRNA-HCR-target_27 | ggggagcaatgatcttgatc**AAA**GCTACCACCACGACATGTACT | 978 | 50.00% |
| BHK-21_ actb-mRNA-HCR-target_28 | cgatccacacagagtacttg**AAA**GCTACCACCACGACATGTACT | 1005 | 50.00% |
| BHK-21_ actb-mRNA-HCR-target_29 | tgatccacatctgctggaag**AAA**GCTACCACCACGACATGTACT | 1053 | 50.00% |
| BHK-21_ actb-mRNA-HCR-target_30 | tagaagcatttgcggtggac**AAA**GCTACCACCACGACATGTACT | 1108 | 50.00% |

| Positive-Control (S10_Cy3+Cy5) | Probe (5'-> 3') | Probe position | Percent GC |
| --- | --- | --- | --- |
| RNA-HCR-Probe-H1_5'-cy5 | cy5 - 5' - TGTTGCAAAGGAACGTCGAGC***TGTAATGGT***GCTCGACGTTCC - 3' |  |  |
| RNA-HCR-Probe-H2_3'-cy5 | 5' - GCTCGACGTTCC***TTTGCAACA***GGAACGTCGAGCACCATTACA - 3' - cy5 |  |  |
| BTV1-WT-S10_RNA-HCR-target_1 | atggcagcgacacttttt***AAA***GCTCGACGTTCCTTTGCAACA | 4 | 44.00% |
| BTV1-WT-S10_RNA-HCR-target_2 | tttggatcagcccggata***AAA***GCTCGACGTTCCTTTGCAACA | 24 | 50.00% |
| BTV1-WT-S10_RNA-HCR-target_3 | tcattttttcttcctcga***AAA***GCTCGACGTTCCTTTGCAACA | 45 | 33.00% |
| BTV1-WT-S10_RNA-HCR-target_4 | ttcaacccgttcctgatt***AAA***GCTCGACGTTCCTTTGCAACA | 68 | 44.00% |
| BTV1-WT-S10_RNA-HCR-target_5 | acacggaccagactcagc***AAA***GCTCGACGTTCCTTTGCAACA | 88 | 61.00% |
| BTV1-WT-S10_RNA-HCR-target_6 | gcggttgggaaatcgtgt***AAA***GCTCGACGTTCCTTTGCAACA | 111 | 56.00% |
| BTV1-WT-S10_RNA-HCR-target_7 | cgcactcggagcatatct***AAA***GCTCGACGTTCCTTTGCAACA | 131 | 56.00% |
| BTV1-WT-S10_RNA-HCR-target_8 | aggcatcgacgatggcat***AAA***GCTCGACGTTCCTTTGCAACA | 152 | 56.00% |
| BTV1-WT-S10_RNA-HCR-target_9 | gctttgtccaagatttca***AAA***GCTCGACGTTCCTTTGCAACA | 181 | 39.00% |
| BTV1-WT-S10_RNA-HCR-target_10 | cacccgttgtatttgaca***AAA***GCTCGACGTTCCTTTGCAACA | 201 | 44.00% |
| BTV1-WT-S10_RNA-HCR-target_11 | cgctttttgtgtttgcgt***AAA***GCTCGACGTTCCTTTGCAACA | 221 | 44.00% |
| BTV1-WT-S10_RNA-HCR-target_12 | gatgcgaatgcagccttc***AAA***GCTCGACGTTCCTTTGCAACA | 241 | 56.00% |
| BTV1-WT-S10_RNA-HCR-target_13 | cacgaaacgcttctgcgt***AAA***GCTCGACGTTCCTTTGCAACA | 261 | 56.00% |
| BTV1-WT-S10_RNA-HCR-target_14 | ctgtctcaacctcacgtc***AAA***GCTCGACGTTCCTTTGCAACA | 281 | 56.00% |
| BTV1-WT-S10_RNA-HCR-target_15 | ctttaagcctcctaggtc***AAA***GCTCGACGTTCCTTTGCAACA | 344 | 50.00% |
| Positive-Control (S10_Cy3+Cy5) | Probe (5'-> 3') | Probe position | Percent GC |
| RNA-HCR-Probe-H1_5'-cy3 | cy3 - 5' - AGTACATGTCGTGGTGGTAGC***TTGTATGAA***GCTACCACCACG - 3' |  |  |
| RNA-HCR-Probe-H2_3'-cy3 | 5' - GCTACCACCACG***ACATGTACT***CGTGGTGGTAGCTTCATACAA - 3' - cy3 |  |  |
| BTV1-WT-S10_RNA-HCR-target_16 | aacaaccgccgctatcag***AAA***GCTACCACCACGACATGTACT | 392 | 56.00% |
| BTV1-WT-S10_RNA-HCR-target_17 | agcttaaacgccacgctc***AAA***GCTACCACCACGACATGTACT | 448 | 56.00% |
| BTV1-WT-S10_RNA-HCR-target_18 | ccattgaggtatctcagc***AAA***GCTACCACCACGACATGTACT | 479 | 50.00% |
| BTV1-WT-S10_RNA-HCR-target_19 | cattgggttcagactctt***AAA***GCTACCACCACGACATGTACT | 500 | 44.00% |
| BTV1-WT-S10_RNA-HCR-target_20 | cccaaattcaccacacct***AAA***GCTACCACCACGACATGTACT | 520 | 50.00% |
| BTV1-WT-S10_RNA-HCR-target_21 | gaccatcatcaggaaggt***AAA***GCTACCACCACGACATGTACT | 542 | 50.00% |
| BTV1-WT-S10_RNA-HCR-target_22 | cttctctcgctttttgcg***AAA***GCTACCACCACGACATGTACT | 562 | 50.00% |
| BTV1-WT-S10_RNA-HCR-target_23 | tgtcaatttgctggttca***AAA***GCTACCACCACGACATGTACT | 582 | 39.00% |
| BTV1-WT-S10_RNA-HCR-target_24 | tcttcatcacttccttct***AAA***GCTACCACCACGACATGTACT | 606 | 39.00% |
| BTV1-WT-S10_RNA-HCR-target_25 | tcctcaccgcatcattat***AAA***GCTACCACCACGACATGTACT | 633 | 44.00% |
| BTV1-WT-S10_RNA-HCR-target_26 | ccatctagcgggactgat***AAA***GCTACCACCACGACATGTACT | 670 | 56.00% |
| BTV1-WT-S10_RNA-HCR-target_27 | gccactctacctactgat***AAA***GCTACCACCACGACATGTACT | 711 | 50.00% |
| BTV1-WT-S10_RNA-HCR-target_28 | acacgatgcagacctcgg***AAA***GCTACCACCACGACATGTACT | 732 | 61.00% |
| BTV1-WT-S10_RNA-HCR-target_29 | gtctgcatcgtgagatca***AAA***GCTACCACCACGACATGTACT | 758 | 50.00% |
| BTV1-WT-S10_RNA-HCR-target_30 | cccgttagacagcagtag***AAA***GCTACCACCACGACATGTACT | 778 | 56.00% |

**Supplementary Table 2. Biotinylated oligos for RNA complex pull-down assay**

| Name of oligo | Sequences | Locations |
| --- | --- | --- |
| S1 bio 300R | 5’ BiosG/AAAAAAAACCTCAAAAGCGTAACTTTGG | S1 nt300 |
| S1 bio 2700R | 5’ BiosG/AAAAAAACTTTCTGTCGCGATACATC | S1 nt2700 |
| S1 bio 3700R | 5’ BiosG/AAAAAAACTTCCGTTTTCGTCAACTG | S1 nt3700 |
| S6 bio 300R | 5’ BiosG/AAAAAAAAGCTTTTGAAGCATTGGAG | S6 nt300 |
| S6 bio 850R | 5’ BiosG/AAAAAAAAATTCGCCCGTTGGAACCC | S6 nt850 |
| S10 bio 300R | 5’ BiosG/AAAAAAAAATCTGTCTCAACCTCACGTC | S10 nt300 |
| S10 bio 700R | 5’ BiosG/AAAAAAAATGGTAATTCGAAACCATCTAG | S10 nt700 |

**Supplementary Table 3. qPCR primers for BTV ten segments**

| **qPCR primers** | **Sequences** | **Size (nt)** | **GC (%)** | **TM (°C)** | **Target size (bp)** |
| --- | --- | --- | --- | --- | --- |
| BTV1-wt_S1-F | 5' TCGACACGCACCTTTCCGG 3' | 19 | 63.16 | 62.9 | 109 |
| BTV1-wt_S1-R | 5' ATCCCTGGCTGCTCCTTCC 3' | 19 | 63.16 | 61.7 |  |
| BTV1-wt_S2-F | 5' ACATTTCATGCGGCGCAGGA 3' | 20 | 55 | 61.3 | 109 |
| BTV1-wt_S2-R | 5' TCTGATCCCCTGGCCTAAACG 3' | 21 | 57 | 64.1 |  |
| BTV1-wt_S3-F | 5' GCCGACCCAGTCGTGCTAG 3' | 19 | 68.42 | 62.5 | 109 |
| BTV1-wt_S3-R | 5' CGATCAGCGGAGCAATCTCG 3' | 20 | 60 | 60.2 |  |
| BTV1-wt_S4-F | 5' CCGTCTCGTTAAAGGAGCCG 3' | 20 | 60 | 60.6 | 109 |
| BTV1-wt_S4-R | 5' GTTGGGTTTCGGACCCTCA 3' | 19 | 57.89 | 60.7 |  |
| BTV1-wt_S5-F | 5' ATGCGGAGACGGTCATGGTG 3' | 20 | 60 | 61.8 | 109 |
| BTV1-wt_S5-R | 5' GCGATCTCCTGTATCGCCTCC 3' | 21 | 61.9 | 64.3 |  |
| BTV1-wt_S6-F | 5' GGATATGCGCCGCGTGCAG 3' | 19 | 68.42 | 63 | 109 |
| BTV1-wt_S6-R | 5' CCCTCTCGTCCTCATCCTCG 3' | 20 | 65 | 62 |  |
| BTV1-wt_S7-F | 5' ACCAGCGCGTCAGCCCTAT 3' | 19 | 63.16 | 62.4 | 109 |
| BTV1-wt_S7-R | 5' CGGACCACACACTACCGCA 3' | 19 | 63.16 | 61.9 |  |
| BTV1-wt_S8-F | 5' CGCCAAGGGAAGAGTCACGC 3' | 20 | 65 | 63 | 109 |
| BTV1-wt_S8-R | 5' CTCGCTTCACGCAGCTTCTC 3' | 20 | 60 | 60.2 |  |
| BTV1-wt_S9-F | 5' CTGCTGAGAGAGGGAGGCG 3' | 19 | 68.42 | 62.2 | 109 |
| BTV1-wt_S9-R | 5' GTGATCCCGACACTCGCTGG 3' | 20 | 65 | 62.8 |  |
| BTV1-wt_S10-F | 5' GCGCCTATGCCATCGTCG 3' | 18 | 66.67 | 60.6 | 110 |
| BTV1-wt_S10-R | 5' GCGAATGCAGCCTTCTCCG 3' | 19 | 63.16 | 61 |  |
